# Regulation of Microprocessor assembly and localization via Pasha’s WW domain in *C. elegans*

**DOI:** 10.1101/2024.04.23.590772

**Authors:** Brooke E. Montgomery, Thiago L. Knittel, Kailee J. Reed, Madeleine C. Chong, Ida J. Isolehto, Erin R. Cafferty, Margaret J. Smith, Reese A. Sprister, Colin N. Magelky, Hataichanok Scherman, Rene F. Ketting, Taiowa A. Montgomery

## Abstract

Primary microRNA (pri-miRNA) transcripts are processed by the Microprocessor, a protein complex that includes the ribonuclease Drosha and its RNA binding partner DGCR8/Pasha. We developed a live, whole animal, fluorescence-based sensor that reliably monitors pri-miRNA processing with high sensitivity in *C. elegans*. Through a forward genetic selection for alleles that desilence the sensor, we identified a mutation in the conserved G residue adjacent to the namesake W residue of Pasha’s WW domain. Using genome editing we also mutated the W residue and reveal that both the G and W residue are required for dimerization of Pasha and proper assembly of the Microprocessor. Surprisingly, we find that the WW domain also facilitates nuclear localization of Pasha, which in turn promotes nuclear import or retention of Drosha. Furthermore, depletion of Pasha or Drosha causes both components of the Microprocessor to mislocalize to the cytoplasm. Thus, Pasha and Drosha mutually regulate each other’s spatial expression in *C. elegans*.

## INTRODUCTION

miRNAs are implicated in nearly every biological process and their dysregulation commonly leads to developmental abnormalities and disease^1^. miRNA biogenesis involves two sequential RNA cleavage steps that release the mature miRNA duplex from the primary miRNA (pri-miRNA) transcript^2^. The first cleavage event occurs within the nucleus where the Microprocessor complex excises the miRNA hairpin from the pri-miRNA to form what is called the pre-miRNA^3–8^. Following transport to the cytoplasm, Dicer removes the terminal loop from the pre-miRNA hairpin to free the miRNA duplex, marking the second cleavage event^9–12^. The miRNA duplex then binds an Argonaute protein, which eliminates one strand and retains the other to serve as a guide for sequence-specific mRNA silencing^9,13^.

Elegant biochemical and structural studies have informed much of our understanding of pri-miRNA processing. The Microprocessor, a complex comprising at least two core proteins - the RNA-binding protein DGCR8/Pasha and the ribonuclease Drosha - identifies pri-miRNA transcripts and initiates miRNA biogenesis^3–5^. Drosha resides at the base of the hairpin, cleaving it to release it from the primary transcript^3,14^. DGCR8/Pasha forms a dimer at the top of the miRNA hairpin, playing a crucial, albeit indirect, role in cleavage of the hairpin^6,14^. The precise function of DGCR8/Pasha remains somewhat uncertain. A short peptide of DGCR8/Pasha, encompassing its Drosha interaction domain, when complexed with Drosha is adequate for cleavage of certain miRNA hairpins. However, longer fragments of DGCR8/Pasha containing the double-stranded RNA (dsRNA) binding domains lead to more effective miRNA processing^14^. Consequently, it is probable that DGCR8/Pasha functions both in dsRNA binding, thereby stabilizing the Microprocessor-pri-miRNA complex, and in ensuring proper folding and orientation of Drosha. Dimerization of DGCR8/Pasha is also important for accurate processing of pri-miRNA transcripts, underscoring its significance in correctly orienting Drosha on the miRNA hairpin^14^. Interestingly, in *C. elegans*, but seemingly not in humans, Pasha also serves as a ruler to guide Drosha cleavage at the correct position relative to the top of the hairpin^15^.

While *in vitro* studies play a crucial role in elucidating the mechanisms of pri-miRNA processing, *in vivo* investigations are essential for understanding the dynamic aspects of this process within the broader cellular context. Therefore, we developed a reporter system for pri-miRNA processing in a genetically tractable whole-animal model to explore the first step in miRNA biogenesis. This system comprises a fluorescent sensor that faithfully reflects the cleavage of a pri-miRNA transcript in *C. elegans*. Utilizing the sensor in a forward genetic screen, we identified a mutation in the G residue of the WW domain of DGCR8/Pasha, a region embedded within the RHED domain crucial for heme binding and protein dimerization^16–18^. Intriguingly, animals with this mutation, or with an engineered mutation in the adjacent namesake W residue of the WW domain, are viable despite a modest but widespread decrease in miRNA levels. Pasha dimerization and Microprocessor integrity is impaired in the G and W mutants, which likely underlies the strong reduction in nuclear enrichment of both Pasha and Drosha in these mutants.

Knockdown of Pasha via RNA interference also disrupts Drosha’s nuclear enrichment, while Drosha knockdown leads to a reduction in Pasha’s nuclear localization. Thus, correct assembly of the Microprocessor is important for nuclear import or retention of both Pasha and Drosha. Furthermore, our findings reveal that while a reduction in the nuclear localization of the Microprocessor is tolerable, exclusion of Pasha from the nucleus results in embryonic or early larval arrest, underscoring a critical role for nuclear activity of the Microprocessor in development.

## RESULTS

### A sensor for primary miRNA processing in *C. elegans*

We developed a sensor to explore the requirements for primary miRNA recognition and processing in live *C. elegans*. The sensor contains the hairpin and regulatory sequences for miR-58, a presumed ortholog of bantam in flies^19^, fused to mCherry coding sequence and expressed under the control of the ubiquitin, *ubl-1*, promoter (Fig. 1A). When the miR-58 hairpin is recognized by the Microprocessor machinery and cleaved, the mCherry mRNA is detached from the *mir-58* 3’ regulatory elements, which would presumably lead to its degradation because Microprocessor cleavage of exonic hairpins leads to mRNA destabilization^20^. Thus, we predicted that the sensor would produce elevated levels of mCherry if pri-miRNA recognition or processing was impaired. To test this, we made a transgenic line in which we integrated the sensor into the genome as a single-copy transgene^21^. We also made a control line with an identical transgene at the same genomic position, but which contained the *ubl-1* 3’UTR in place of the *mir-58* sequence. In the control strain that lacked *mir-58* sequence, mCherry was strongly and similarly expressed in animals treated with vector control or *pash-1* RNAi (Fig. 1B). In contrast, the pri-miR-58 sensor was weakly expressed on control RNAi treatment but was strongly expressed on *pash-1* RNAi (Fig. 1B). *pash-1* and *drsh-1* are both essential but first-generation homozygous animals are viable^5^. Thus, we introduced the sensor into *drsh-1* mutants, homozygosed the sensor, and then imaged animals segregating *drsh-1+/+* and *drsh-1-/-* in the first generation of homozygosity in which mutant animals are still healthy. The sensor was strongly desilenced in *drsh-1-/-* mutant relative to *drsh-1+/+* wild type animals (Fig. 1B). Therefore, the sensor is responsive to loss of both *pash-1* and *drsh-1*, the core components of the Microprocessor.

**Figure 1.**
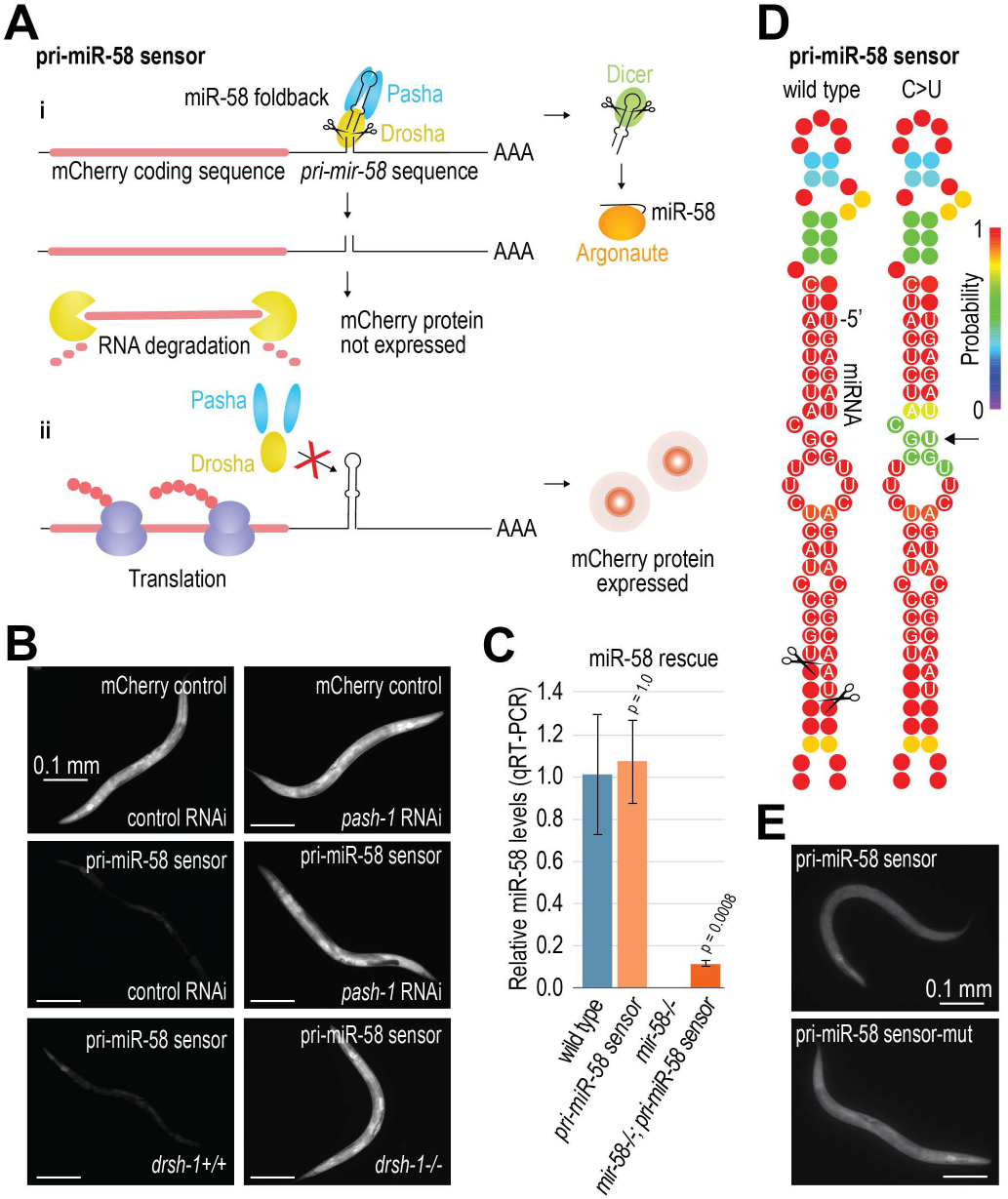
A sensor for pri-miRNA recognition and processing in *C. elegans.* (A) pri-miR-58 sensor design. i) In the presence of a functional miRNA biogenesis pathway, the miR-58 hairpin is cleaved from the mCherry mRNA and routed through the miRNA pathway, leading to the degradation of mCherry. ii) If the sensor is not recognized and processed, mCherry is expressed. (B) mCherry fluorescence in animals transformed with the pri-miR-58 sensor or a control construct lacking pri-miR-58 sequence. Animals were treated with either control (empty L4440 vector) or *pash-1* RNAi, or were segregants for wild type *drsh-1* or the *drsh-1* deletion allele *drsh-1(ok369).* (C) Relative levels of mature miR-58 levels in the various strains indicated as determined by TaqMan qRT-PCR. Error bars are standard deviation from the mean 3 biological replicates. P values are for comparison to wild type. (D) Secondary structure predictions of the wild type and C-U mutant miR-58 hairpin. Nucleotide sequence is shown for the miRNA duplex region. The 5’ end of the miRNA is indicated. (E) mCherry fluorescence in animals containing the unaltered or C-U mutant pri-miR-58 sensor construct.

The pri-miR-58 sensor differs from typical endogenous pri-miRNAs in that it contains protein coding sequence. This mCherry coding sequence could interfere with the usual processing of the precursor through the miRNA biogenesis pathway. To assess whether the sensor generated mature miR-58, indicating effective processing through the miRNA pathway, we evaluated its capability to restore mature miR-58 levels in a *mir-58* knockout mutant^22^. The sensor restored miR-58 levels in *mir-58*-/- mutants to ∼11% of wild type levels but had a negligible impact on miR-58 levels in *mir-58*+/+ animals (Fig. 1C). Hence, the sensor generates mature miR-58, albeit at levels nearly 10-fold lower than those from the endogenous locus. This reduction may be attributed to the translational machinery sequestering the transcript away from the miRNA pathway due to the presence of the mCherry coding sequence. In support of this possibility, the reporter still produced mCherry protein, although at relatively low levels, in otherwise wild type animals, suggesting competition between the miRNA biogenesis and mRNA translation pathways for the reporter transcript (Fig. 1B). This sensor has the advantage over other sensors^23^ in that it reports on pri-miRNA processing in whole live animals and can be scored on a standard stereo microscope. As such, we believe that it will be a valuable resource to the small RNA community.

### An expanded bulge inhibits pri-miR-58 processing

We then designed a forward genetic screen to identify nucleotides within or near the miR-58 hairpin important for recognition and processing by the Microprocessor. Heterozygous cis-acting mutations that disrupt cleavage of one copy of the sensor should partially desilence mCherry expression, whereas only dominant or dosage-dependent heterozygous mutations in trans-acting factors would lead to desilencing. Therefore, we enriched for mutations in the sensor over mutations in trans-acting genes by screening the first-generation progeny of mutagenized animals, which should be heterozygous for any mutations. We identified 500 animals with bright mCherry fluorescence and then sequenced the miR-58 hairpin from each of these individuals. We identified a single mutation in the hairpin, positioned within the miRNA duplex region (Fig. 1D). The mutation changes a G-C base pair to a G-U base pair adjacent to the central bulge in the hairpin. The mutation likely causes the bulge to expand, thereby preventing recognition or binding of the Microprocessor and thus interfering with cleavage. The site of the mutation is not conserved amongst *C. elegans* miRNAs suggesting that it is not part of a sequence motif important for miRNA processing (Supplemental Fig. S1A).

Because we sacrificed this mutant animal to identify the causal mutation, we engineered the mutation into the sensor to confirm that it was responsible for mCherry desilencing. Indeed, a modest increase in mCherry fluorescence was observed in the mutant relative to the original form of the sensor (Fig. 1E). We draw two conclusions from these results. Firstly, the recognition or processing of the miR-58 hairpin is susceptible to slight perturbations in the miRNA duplex region. Secondly, the sensor accurately reflects the recognition and processing of the miR-58 hairpin by the Microprocessor machinery.

### A GW motif within Pasha’s WW domain promotes cleavage of pri-miR-58

Having confirmed that the pri-miR-58 sensor provides a reliable reporter for recognition and processing by the Microprocessor, we then did a forward genetic selection for alleles that desilenced mCherry. This time, we allowed two generations for the mutagenized animals to become homozygous for mutant alleles during the selection process to identify trans-acting factors. We identified 72 independent lines that desilenced mCherry, only 17 of which were fertile over multiple generations. Of the 17 that were fertile, only one line, 38a, displayed a strong increase in mCherry levels (Fig. 2A-2B). We subjected this line to whole genome sequencing and identified a missense mutation in *pash-1* (Fig. 2C). We backcrossed the line to the original non-mutated line and confirmed that mCherry desilencing tracked perfectly with the *pash-1* allele, indicating that it was likely the causal mutation. We then outcrossed the allele, *pash-1(ram33)*, to wild type animals three times to remove the sensor and clear some of the background mutations.

**Figure 2.**
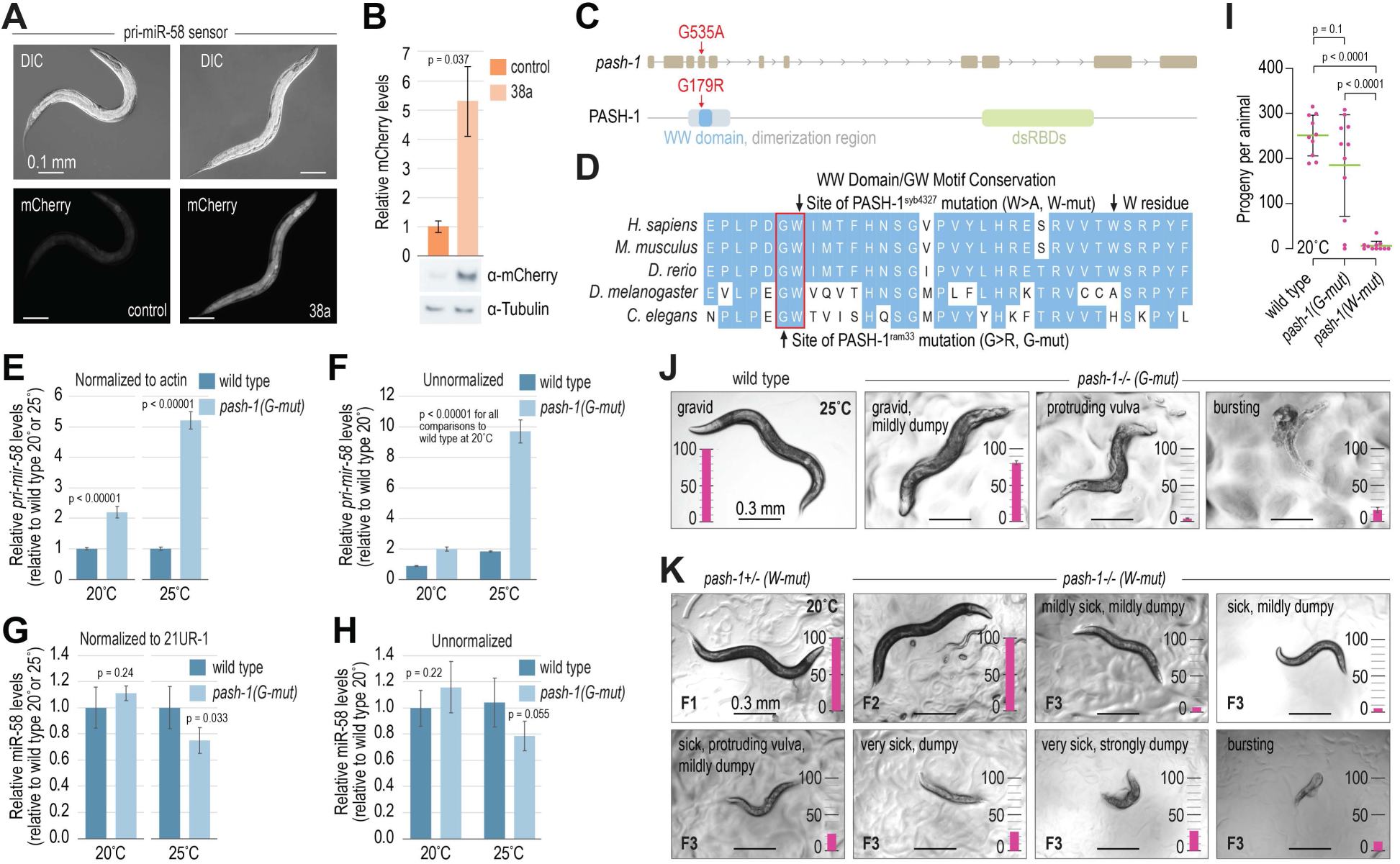
Pasha’s WW domain is required for pri-miRNA recognition or processing. **(A)** Representative images showing mCherry fluorescence in control (non-mutated) and 38a-mutant animals containing the pri-miR-58 sensor construct. DIC and mCherry fluorescence images are shown. (B) Relative mCherry levels in control and 38a-mutant animals containing the pri-miR-58 sensor construct as determined by Western blot analysis (the blot image of 1 of 3 biological replicates is shown below the plot). Tubulin was used for normalization. Error bars are standard deviation from the mean of 3 biological replicates. (C) The location of the 38a mutation on *pash-1* DNA and protein sequences. The WW, dimerization, and dsRNA-binding domains are shown. (D) Sequence and conservation of the WW domain. The 38a *(pash-1(ram33), G-mut)* and *pash-1(syb4327) (W-mut)* mutants are indicated. (E-F) Relative levels of endogenous pri-miR-58 in wild type and *pash-1(ram33)* mutants grown at 20^°^C or 25^°^C as determined TaqMan qRT-PCR. Data is normalized to *act-1* mRNA levels (E) or not normalized (F). Error bars are standard deviation from the mean of 3 biological replicates. P values are for comparison to wild type. (G-H) Relative levels of mature miR-58 in wild type and *pash-1(ram33) (G-mut)* mutants grown at 20^°^C or 25^°^C as determined TaqMan qRT-PCR. Data is normalized to 21 UR-1 piRNA levels (G) or not normalized (H). Error bars are standard deviation from the mean 3 biological replicates. P values are for comparison to wild type. (I) Numbers of progeny produced by wild type and *pash-1(ram33) (G-mut)* and *pash-1(syb4327) (W-mut)* animals grown at 20^°^C. Error bars are standard deviation from the mean for 10-11 animals. (J-K) Representative images of wild type and *pash-1(ram33) (G-mut)* (J) or *pash-1(syb4327) (W-mut)* (K) animals grown at 20^°^C or 25^°^C as indicated. The bar plots show the percentage of animals with the phenotype indicated (n = 100 animals per strain).

The *pash-1(ram33)* mutation changes a glycine (G) residue within Pasha’s WW domain to an arginine (R) (Fig. 2C). This residue is adjacent to the first tryptophan (W) residue of the WW domain which is embedded in the broader RHED domain that binds heme in humans. It is important to note that in *C. elegans* and *D. melanogaster*, Pasha lacks the second W residue found in most WW domains^24^. The G residue that is mutated in *pash-1(ram33)* is highly conserved in DGCR8/Pasha and other WW domain-containing proteins (Fig. 2D)^24^. We will hereafter refer to the *pash-1(ram33)* allele as the G mutant (G-mut) for simplicity and to distinguish it from another mutation we describe below.

We then assessed by qRT-PCR the impact of the G mutation on the endogenous *pri-mir-58* transcript in *pash-1(ram33)* after having removed the sensor. When grown at 20°C, levels of *pri-mir-58* were upregulated ∼2 fold in animals containing the G mutation compared to wild type (Fig. 2E). At 25°C, a less favorable growth temperature that often enhances molecular and development defects in *C. elegans*, *pri-mir-58* levels were elevated ∼5-fold in G-mutant animals, indicating the allele is somewhat temperature sensitive (Fig. 2E). We could not directly compare *pri-mir-58* levels at 20°C and 25°C because of possible differences in expression of the housekeeping genes used for normalization at the two temperatures. However, we reanalyzed the data without normalization to the housekeeping genes and observed nearly identical fold changes between G-mutant and wild type animals compared to what was observed after normalization to actin, indicating that a direct comparison without normalization is reasonable (Fig. 2F). From this comparison, we identified an ∼2-fold increase in *pri-mir-58* levels at 25°C relative to 20°C, suggesting that either *mir-58* transcription is upregulated or that *pri-mir-58* processing is less efficient at 25°C (Fig. 2F). If pri-miRNA processing is less efficient at 25°C, it could explain why miRNA processing is more severely impaired in animals harboring the G-mutant at this temperature. Interestingly, mature miR-58 levels were essentially unchanged at 20°C and reduced by only ∼25% at 25°C in G-mutant animals relative to wild type (Fig. 2G). Analyzing the data without normalization to an unrelated small RNA (the piRNA 21UR-1) did not change these results, but unlike the primary transcript, mature miR-58 levels were unchanged at 25° relative to 20°C in wild type animals (Fig. 2H). The lack of correlation we observed between primary and mature miR-58 suggests that the amount of pri-miR-58 exceeds what can be processed by the miRNA machinery. More conclusively, however, these results reveal a modest and temperature-sensitive impact of Pasha’s G residue on endogenous pri-miRNA processing.

### Mutations in the WW domain cause a spectrum of developmental defects

Animals harboring the Pasha G-mutant are viable, and when grown at 20°C tended to produce nearly as many progeny as wild type (Fig. 2I). However, there was a higher proportion of animals producing very few progeny which led to an overall decline in mean brood size (Fig. 2I). When grown at 25°C, G-mutant animals consistently produced fewer progeny than wild type and ∼50% were sterile (Supplemental Fig. S1B). Nonetheless, even when grown at 25°C, most animals appeared remarkably healthy aside from a slight squatty (i.e. dumpy) phenotype, although some had protruding vulvas or extrusion of their guts through their vulvas (i.e. bursting), which are phenotypes common to *C. elegans* miRNA mutants (Fig. 2J)^9^.

Attempts to recover mutations in the WW domain of Pasha in flies have been unsuccessful, likely because such mutations cause lethality^25^. Hence, it was surprising to us that G-mutant animals were relatively healthy. Given the proximity of the G residue to the W residue of the WW domain, it is possible that the G mutation causes only modest perturbation to the domain’s function. To test this possibility, we mutated the W residue to alanine (A) using CRISPR-Cas9 genome editing (Fig. 2D)^26–29^. We refer to this mutant as the W mutant (W-mut). The pri-miR-58 sensor was desilenced when introduced into W-mutant animals, indicating that the W residue is required for efficient pri-miRNA processing (Supplemental Fig. S1C). Unlike G-mutants, W-mutant animals were nearly sterile at 20°C and completely sterile at 25°C (Fig. 2I and Supplemental Fig. S1B). W-mutants also displayed a spectrum of developmental defects, including a range in dumpy, protruding vulva, and bursting phenotypes (Fig. 2K). These defects were absent in the first generation of homozygosity of the W mutation (F2). Instead, first generation homozygous W-mutants were superficially wild type, suggesting that a maternal contribution of wild type Pasha is sufficient for normal development and indicating that the mutation is not dominant (Fig. 2K). Thus, the W residue, while not critical for viability, is important for normal development. Although the W residue is a defining feature of WW domains, we questioned whether the G and W residues, collectively referred to as the GW motif, have distinct or additive functionality. To address this question, we generated a strain containing both the G>R and W>A substitutions. Surprisingly, animals containing both mutations produced significantly fewer progeny than either of the single mutants (p = 0.015) (Supplemental Fig. S1D; note that these mutations were made in the *pash-1::GFP* background described below). Thus, the G and W residues likely have independent contributions to the function of this domain.

### Widespread loss of miRNAs in WW domain mutants

To identify the impact of the G mutation on mature miRNA levels, we subjected wild type and G-mutant adult animals grown at either 20°C or 25°C to small RNA high-throughput sequencing. Most canonical miRNAs were modestly but significantly downregulated in the G-mutant animals relative to wild type when grown at 20°C (Fig. 3A; Supplemental Data 1). At 25°C, a less ideal growth condition for *C. elegans*, we observed a more substantial reduction in miRNA levels in G-mutant animals, however, because the animals are mildly sick at this temperature, developmental differences limit the interpretation of individual miRNA levels (Fig. 3B; Supplemental Data 1). A small number of miRNAs were upregulated in the G-mutant animals at both 20°C and 25°C, most notably mirtrons, which do not require the Microprocessor for their biogenesis (Fig. 3C; Supplemental Data 1)^30,31^. Additionally, the miR-1829 family comprising 4 miRNAs and the miR-42 cluster consisting of 2 miRNAs were also upregulated, which could indicate that these miRNAs are also not dependent on the Microprocessor for their biogenesis (Fig. 3C; Supplemental Data 1). We did not confirm that the miRNAs identified in Ahmed et al. are authentic miRNAs, but it is possible that these are also non-Microprocessor dependent miRNAs as they were also upregualted^32^. It is possible that in G mutants there is reduced competition for shared resources downstream of pri-miRNA processing due to the likely reduction in pre-miRNAs, which are subsequently diverted to mirtrons and other Microprocessor-independent miRNAs.

**Figure 3.**
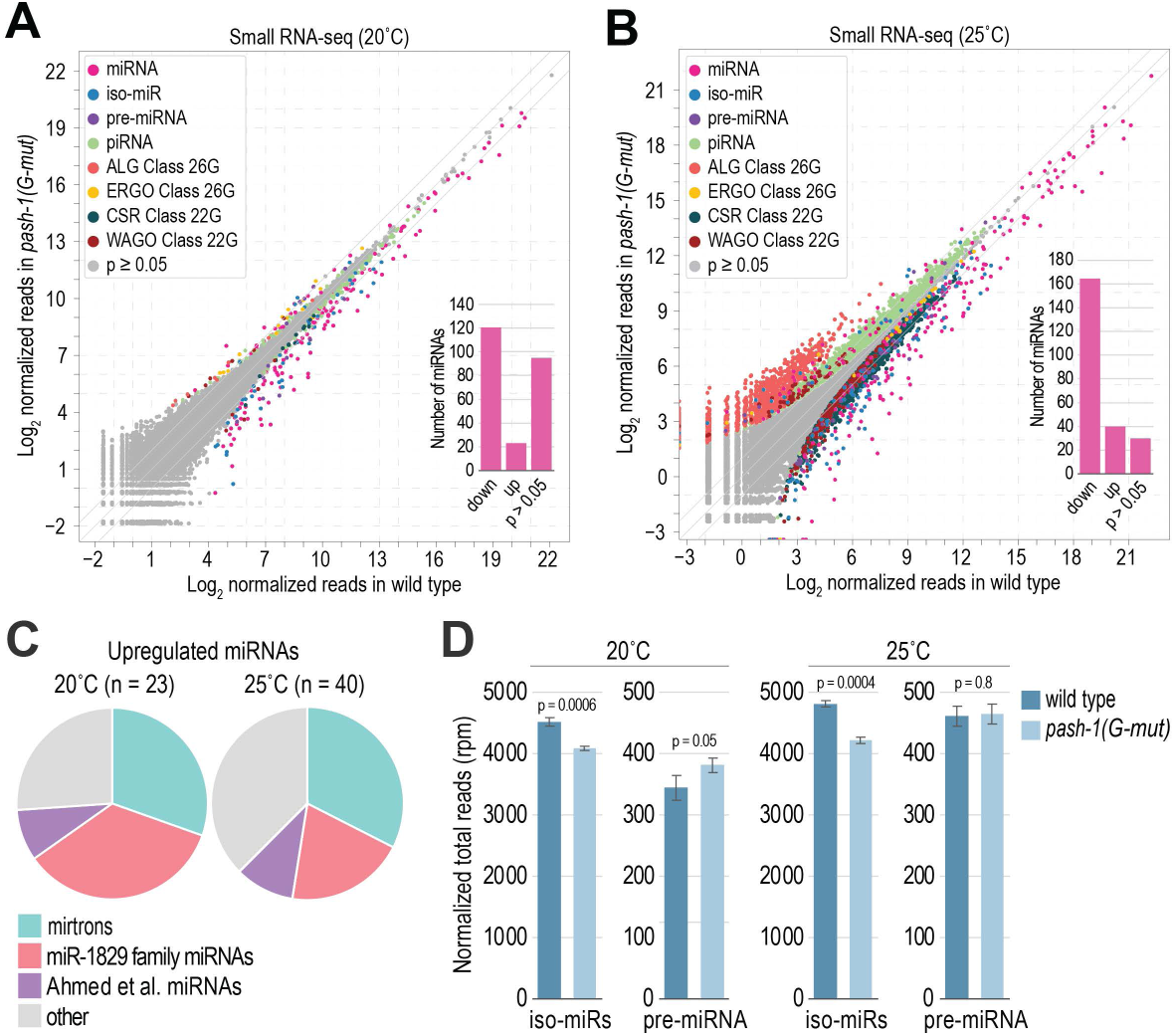
Widespread reduction in canonical miRNA levels in Pasha G-mutants. (A-B) Log**_2_** average geometric-mean normalized small RNA read counts in wild type (x-axis) and *pash-1(ram33)* (G-mut) (y-axis) animals grown at 20^°^C (A) or 25^°^C (B) as determined by sRNA-seq_ Small RNAs are represented by data points colored by their classification. The inset bar plots show the numbers of miRNAs represented by >20 geometric mean normalized reads significantly down or upregulated (p < 0.05) or unchanged in *pash-1(ram33)* relative to wild type. Data is from 3 (A) or 4 (B) biological replicates per strain. (C) Classification of miRNAs upregulated in *pash-1(ram33)* relative to wild type animals grown at either 20^°^C or 25^°^C, as indicated, based on data in (A) and (B)_ (D) Mean reads per million (rpm}-normalized counts for iso-miRs (offset by 1-3 nt relative to miRNA 5’ end) and pre-miRNA (pre-miRNA derived reads offset by >3 nt relative to miRNA 5’ end) in wild type and *pash-1(ram33)* mutants grown at 20^°^C or 25^°^C. Data as in (A) and (B). Error bars are standard deviation from the mean for 3 (20^°^C} or 4 (25^°^C} biological replicates. P values are for comparison to wild type.

Dimerization of Pasha/DGCR8 via the RHED domain region, which contains the WW domain, was proposed to have a role in miRNA processing precision based on imprecise processing of miRNA hairpins by Microprocessor complexes containing only one molecule of human DGCR8^14^. Thus, the reduction in mature miRNA levels we observed in *pash-1* G-mutants could be due to misprocessing, rather than loss of processing, of pri-miRNAs. However, we did not observe an increase in individual miRNA isoforms (iso-miRs) shifted by 1-3 nucleotides (nts) at their 5’ ends or in pre-miRNA reads offset by >3 nts in G-mutants as would be expected if processing precision was impaired (Fig. 3A-3B; Supplemental Data 1). In fact, total iso-miR reads were slightly depleted in G-mutants (Fig. 3D). Although there was a slight increase in pre-miRNA reads in G-mutants grown at 20°C, it was diminished at 25°C (Fig. 3D). Furthermore, mutations in both the G and W residues desilence the pri-miR-58 sensor, and because cleavage anywhere within the hairpin of the sensor would presumably cause desilencing, this reflects a loss of processing, rather than a change in processing precision (Fig. 2A and Supplemental Fig. S2B). Together, these results suggest that it is unlikely that the G and W residues, and by extension the WW domain, contribute substantially to pri-miRNA processing precision in *C. elegan*s.

Severe developmental defects and inconstancies in developmental progression precluded meaningful analysis of global miRNA levels in W-mutant animals. However, through a very limited analysis of miR-1, let-7, and miR-58 by qRT-PCR we confirmed that these miRNAs are modestly depleted in W-mutants, just as in G-mutants, with the caveat that we did not control for animal health and morphology (Supplemental Fig. S2). The modest depletion observed here is consistent with what was observed in HeLa cells containing a W>A mutation at the second W residue of DGCR8’s WW domain (which is not conserved in worms, as noted above) and point to a similar requirement for the WW domain in miRNA processing in humans and nematodes^23^.

We conclude that Pasha’s G and W residues are necessary for optimal miRNA biogenesis, although they are not essential. Additionally, the relatively good health of G mutants and the viability of W mutants, despite significantly lower miRNA levels, indicate that *C. elegans* has a high tolerance for reduced miRNA activity. This tolerance might explain why the loss of most individual miRNAs does not result in severe developmental defects^22^.

### Pasha and Drosha localize throughout the cell

To investigate the subcellular localization of the *C. elegans* Microprocessor and to determine if it varies in Pasha G and W mutants, we used CRISPR-Cas9 to introduce GFP as a C-terminal fusion with Pasha and mCherry as an N-terminal fusion with Drosha at their respective genomic locations^26–29^. We verified that the insertion of GFP and mCherry did not interfere with miRNA biogenesis, as mature miRNA levels remained unchanged in these strains (Supplemental Fig. S3A). Both Pasha and Drosha fusion proteins localized to the nucleus, as previously observed in human cells, with relatively strong expression in the germline and throughout embryogenesis (Fig. 4A-4B and Supplemental Fig. S3B)^33–35^. Consistent with observations in human cells^33,36^, we also observed nucleolar enrichment of Pasha and Drosha, particularly in oocytes, although there was considerable variation between animals (Fig. 4A-4B). Pasha and Drosha co-localized with each other and promptly reentered the nucleus after cell division, appearing to do so simultaneously (Supplemental Fig. S3C). However, they occasionally formed distinct foci within the nucleus (Supplemental Fig. S3D). We observed that while Pasha and Drosha were predominantly localized in the nucleus, they also had a noticeable cytoplasmic signal, although, the signal from PASH-1::GFP was somewhat masked by background fluorescence (Fig. 4A and 4C and Supplemental Fig. S3B-S3C). Based on planar fluorescence quantification, the average signal of mCherry::DRSH-1 was ∼2-4 fold greater in the nucleus compared to the cytoplasm, which varied somewhat based on the developmental stage (Fig. 4C-4D). These results demonstrate that nuclear enrichment of the Microprocessor is conserved in *C. elegans* but point to the existence of both nuclear and cytoplasmic fractions of Pasha and Drosha.

**Figure 4.**
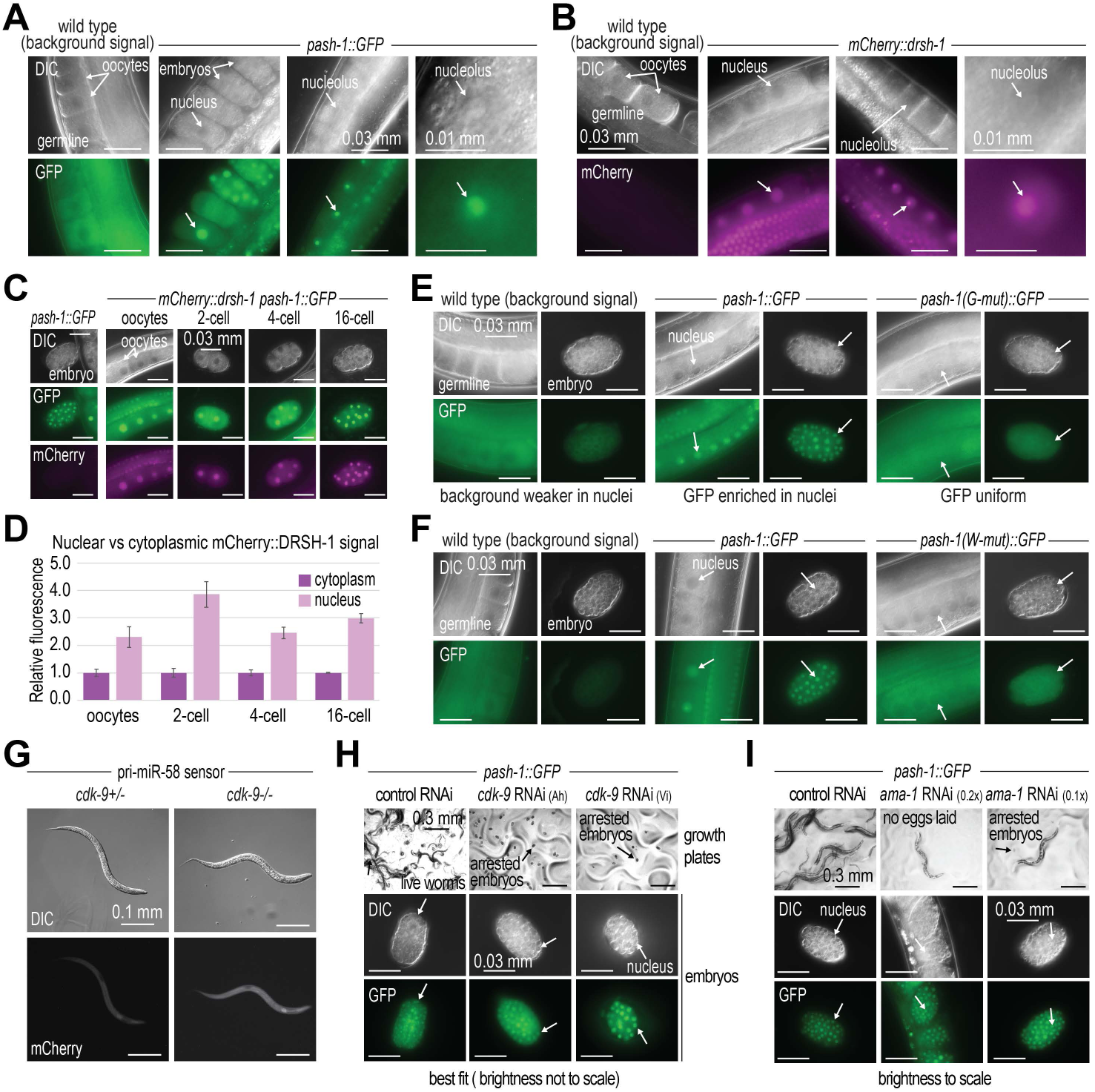
Pasha’s WW domain promotes its nuclear localization. (A) Representative images of PASH-1::GFP expression in the adult germline and embryos *(in utero).* A wild type animal is shown to highlight the non-specific background signal observed in germline tissue. (B) Representative images of mCherry::DRSH-1 expression in the adult germline. A wild type animal is shown to highlight the lack of non-specific background signal observed in germline tissue. (C) Representative images of PASH-1::GFP and mCherry::DRSH-1 expression in oocytes and 2, 4, and 16 cell embryos. A PASH-1::GFP embryo lacking mCherry::DRSH-1 is shown to highlight the lack of non-specific background signal observed for mCherry::DRSH-1 in embryos. Note both nuclear and cytoplasmic signal. (D) Relative mCherry fluorescence signal in the embryos in (C). Error bars are standard deviation from the mean for 2 biological replicates. (E) Representative images of PASH-1 ::GFP, PASH-1 (G-mut)::GFP, and PASH-1 (W-mut)::GFP expression in the adult germline and embryos. A wild type germline and embryo is shown to highlight the non-specific background signal observed in germline tissue. (F) Representative images of PASH-1::GFP and PASH-1 (W-mut)::GFP expression in the adult germline and embryos. (G) Representative images showing mCherry fluorescence in *cdk-9(tm2884)+/­* and *cdk-9(tm2884)-/-* animals containing the pri-miR-58 sensor construct. (H) Representative images showing PASH-1::GFP localization in control (empty L4440 vector) or *cdk-9* RNAi-treated animals. Two different *cdk-9* RNAi clones were used (Ah, Ahringer; Vi, Vidal). Images of growth plates showing embryonic arrest caused by *cdk-9* knockdown. (I) Representative images showing PASH-1::GFP localization in control (L4440) or *ama-1* RNAi-treated animals. Two different concentrations of the *ama-1* RNAi clone were used (0.1x or 0.2x, diluted with L4440). Images of growth plates showing the lack of deposited embryos on the plate (0.2x) or arrested embryos (0.1x) caused by *ama-1* knockdown.

### Pasha mislocalizes to the cytoplasm in WW domain mutants

We then asked if the G and W mutations disrupt subcellular localization of Pasha. We detected PASH-1::GFP, as well as the G- and W-mutant forms, in the nucleus and cytoplasm of cells in germline tissue and embryos, although cytoplasmic expression was somewhat masked by background fluorescence, as noted above (Fig. 4E). Interestingly, the nuclear-to-cytoplasmic signal enrichment observed for PASH-1::GFP was absent in both the G and W mutants (Fig. 4F). Therefore, the G and W residues promote nuclear localization or retention of Pasha. We did not observe a further reduction in nuclear signal in PASH-1(GW-mut)::GFP, which contains both the G and W mutations, indicating that there is not additive effect of the two mutations (Supplemental Fig. S4A).

In *Drosophila*, Pasha’s WW domain promotes association with RNA Polymerase II (Pol II), thereby coupling pri-miRNA transcription to miRNA processing^25^. Hence, Pol II could promote nuclear retention of Pasha in *C. elegans*, which might then explain why its nuclear localization is lost in GW motif mutants. To test this, we first assessed whether deletion of *cdk-9*, a kinase that phosphorylates the C-terminal domain (CTD) of Pol II and which is required for association of Pasha with Pol II in *Drosophila*, desilenced the pri-miR-58 sensor, which would implement it in pri-miRNA processing^25,37^. Although loss of *cdk-9* leads to larval arrest, which could confound the results, we did observe a modest increase in mCherry expression in first generation animals homozygous mutant for *cdk-9*, relative to their heterozygous counterparts, suggesting that it may have a role, albeit possibly indirect, in pri-miRNA processing (Fig. 4G)^38^. We then did RNAi against *cdk-9* in *pash-1::GFP* animals to assess whether loss of *cdk-9* activity affects Pasha’s localization. Neither of two different RNAi clones, both of which caused potent embryonic arrest indicative of *cdk-9* knockdown, disrupted PASH-1::GFP localization, suggesting that phosphorylation of Pol II’s CTD is not required for nuclear retention of the Microprocessor (Fig. 4H).

To assess more directly a requirement of Pol II for Pasha’s localization, we did RNAi against the large subunit of Poll II, *ama-1*^39^. RNAi treatment against *ama-1* using dsRNA-expressing bacteria diluted 5x led to embryonic arrest and retention of embryos *in utero* presumably because of the essential role of Pol II in transcription (Fig. 4I). A 10x dilution of the RNAi treatment (1:10 *ama-1* dsRNA expressing bacteria:vector control bacteria) still caused embryonic arrest but arrested eggs were laid on the plate (Fig. 4I). However, neither treatment caused PASH-1::GFP to mislocalize to the cytoplasm (Fig. 4I). Furthermore, we were not able to detect an interaction between PASH-1::GFP and Pol II by protein co-immunoprecipitation (co-IP) (Supplemental Fig. S4B). Therefore, we conclude that Pasha’s nuclear localization is not facilitated by an association with Pol II and thus that the G and W mutants do not mislocalize due to loss of Pol II association with the Microprocessor.

### Pasha and Drosha crossregulate each other’s localization

To determine if the G and W mutants affect Drosha’s localization, we introduced *mCherry::drsh-1* into strains containing *pash-1(G-Mut)::GFP* or *pash-1(W-Mut)::GFP*. Surprisingly, in both the G and W mutants, mCherry::DRSH-1 also lost its nuclear enrichment (Fig. 5A and Supplemental Fig. S4C-S4F). While imaging these embryos, we noticed a mild and variable reduction in the expression of *pash-1::(G-mut)::GFP* and *pash-1::(W-mut)::GFP*. In some embryos, the cytoplasmic GFP signal from PASH-1(G-mut)::GFP or PASH-1(W-mut)::GFP was stronger than what we typically observed for PASH-1::GFP but in other instances it appeared lower (Supplemental Fig. S4C-S4F). In contrast, mCherry::DRSH-1 was consistently more highly expressed in the cytoplasm of Pasha G and W mutants (Supplemental Fig. S4C-S4F). By Western blot analysis of whole gravid adult animals, we observed ∼50% lower levels of PASH-1::GFP in *pash-1(G-mut)::GFP* and *pash-1(W-mut)::GFP* animals than in *pash-1::GFP* animals but similar levels of mCherry::DRSH-1 (Supplemental Fig. S4G-S4H). Therefore, Pasha’s G and W residues play important roles both in maintaining its stability and in promoting its localization or retention in the nucleus, which in turn is important of proper localization of Drosha.

**Figure 5.**
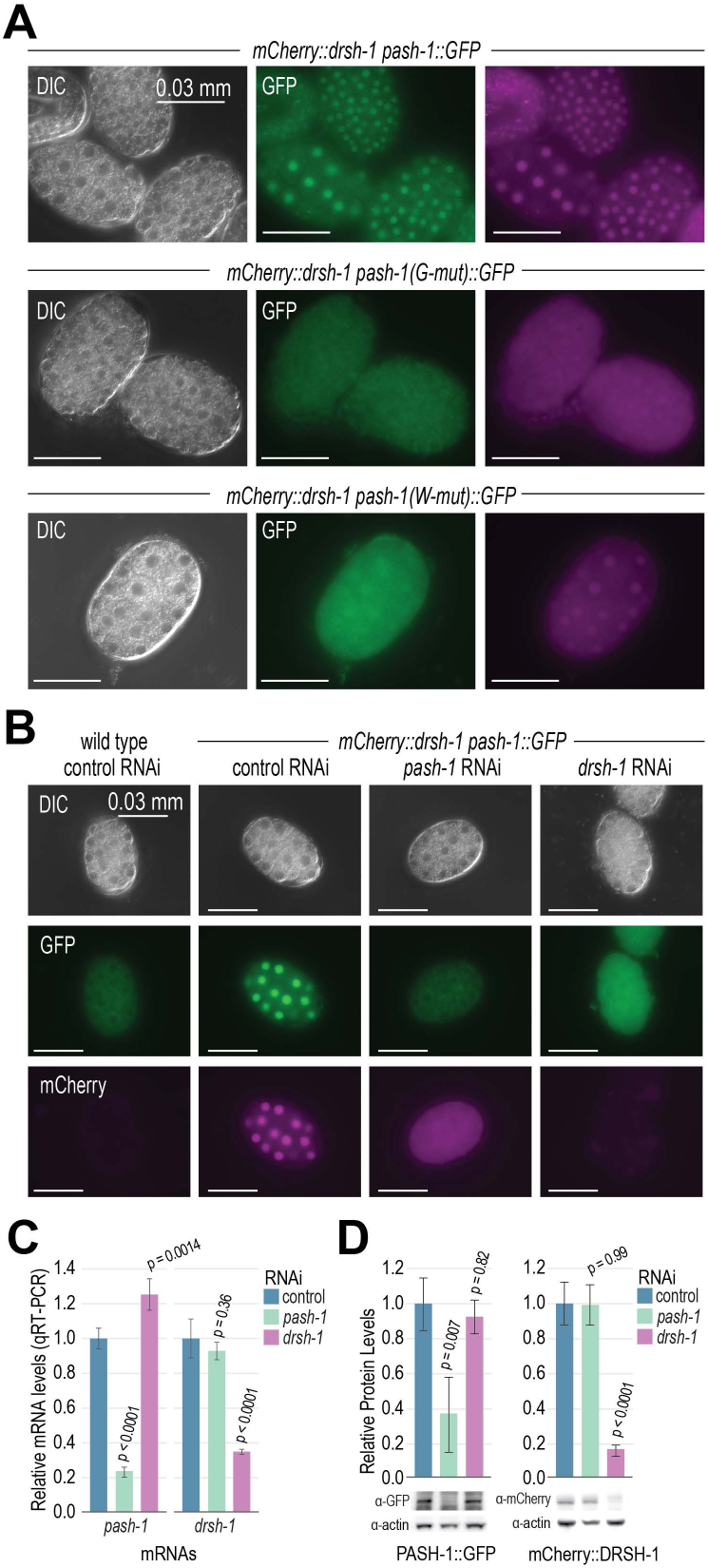
Drosha and Pasha promote each other’s nuclear localization. (A) Representative images of PASH-1 ::GFP, PASH-1 (G-mut)::GFP, PASH-1 (W-mut)::GFP, and mCherry::DRSH-1 expression in embryos. (B) Representative images of PASH-1 ::GFP and mCherry::DRSH-1 expression in embryos following control (L4440), *pash-1,* and *drsh-1* RNAi. (C) Relative *pash-1* and *drsh-1* mRNA levels following control (L4440), *pash-1,* and *drsh-1* RNAi as determined by qRT-PCR. *rpl-32* mRNA levels were used for normalization. Error bars are standard deviation from the mean of 3 biological replicates. P values are for comparison to control RNAi. (D) PASH-1 and DRSH-1 protein levels after *pash-1* or *drsh-1* RNAi. Error bars are standard deviation from the mean of 3 biological replicates. P values are for comparison to control RNAi. One representative of 3 biological replicates is shown in the Western blot. Actin was used for normalization.

The absence of nuclear enrichment of Drosha in Pasha G and W mutants suggests a fundamental role for Pasha in facilitating Drosha’s correct localization. Indeed, depletion of Pasha by RNAi caused mCherry::DRSH-1 to mislocalize to the cytoplasm (Fig. 5B and Supplemental Fig. S5A-S5B). Furthermore, RNAi-knockdown of Drosha caused PASH-1::GFP to lose its nuclear enrichment (Fig. 5B and Supplemental Fig. S5A-S5B). Therefore, both Pasha and Drosha are essential for the correct nuclear localization of the Microprocessor components.

In mammals, the Microprocessor regulates of each of its core constituents, DGCR8/Pasha and Drosha^20^. It downregulates DGCR8/Pasha through cleavage of a hairpin in the 5’ UTR of its mRNA and upregulates Drosha through protein-protein interactions^20^. We took advantage of the Pasha and Drosha RNAi-knockdown assays described above to assess whether this crosstalk is conserved in *C. elegans*. RNAi-knockdown of *pash-1* and *drsh-1* reduced their mRNA levels by ∼75% and ∼65% respectively and led to a consistent decrease in PASH-1::GFP and mCherry::DRSH-1 protein levels, indicating that the RNAi treatment was effective (Fig. 5B and 5D). *drsh-1* knockdown led to an ∼25% increase in *pash-1::GFP* mRNA levels but had no discernable impact on PASH-1::GFP protein levels (Fig. 5C-5D). In *C. elegans*, *pash-1* lacks the hairpin in its 5’ UTR found in other species^20^. Nevertheless, it could still be indirectly regulated by the Microprocessor, as it is predicted to be a target of miR-71. Consistent with this possibility, we previously observed a modest but significant increase in *pash-1* mRNA levels in the absence of the major miRNA Argonaute *alg-1*^40,41^. However, we did not detect an increase in *pash-1* mRNA levels in *mir-71* deletion mutants (Supplemental Fig. S5C). Furthermore, PASH-1::GFP protein levels were not upregulated when we scrambled the *mir-71* binding site, suggesting that the modest increase in *pash-1* mRNA levels following *drsh-1* knockdown may be indirect and inconsequential (Supplemental Fig. S5D).

RNAi knockdown of *pash-1* did not significantly affect *drsh-1* mRNA or protein levels, suggesting that *drsh-1* expression levels are not regulated by the Microprocessor despite its nuclear localization being dependent on Pasha (Fig. 5C-5D). Additionally, RNAi knockdown of the two major miRNA Argonautes, *alg-1* and *alg-2*, did not appear to increase the fluorescence signals from either PASH-1::GFP or mCherry::DRSH-1 in embryos (Supplemental Fig. S5E). Nor did *alg-1* and *alg-2* knockdown affect Pasha and Drosha localization to the nucleus, indicating that miRNAs do not regulate subcellular localization of the Microprocessor (Supplemental Fig. S5E). These results suggest that any crossregulation that might exist between Pasha and Drosha in *C. elegans* is not likely to have a substantial role in their protein expression levels. However, given that both proteins depend on each other for their nuclear localization, they nonetheless share an important functional regulatory interaction.

### Pasha’s WW domain promotes dimerization and Microprocessor assembly

It is possible that the loss of nuclear localization of Pasha and Drosha in the Pasha G and W mutants is due to misassembly of the Microprocessor. To determine if Microprocessor formation is affected by mutations in the GW motif of the WW domain, we first did size exclusion chromatography on animals expressing mCherry fused to Drosha and either wild type or the G or W mutant form of Pasha fused to GFP. We observed several fractions between 158-669 kDa that contained both Drosha and Pasha (Fig. 6A). Western blot signals for PASH-1::GFP and mCherry:DRSH-1 were substantially lower for the Pasha G and W mutants, which may relate to both lower levels of mutant PASH-1::GFP, as noted above, as well as lower solubility of the mutant proteins, as was observed for fly and human WW domain mutants^16,25^. Despite the caveat of reduced protein levels, the large complex still formed in both G and W mutants, suggesting that these residues are not absolutely essential for complex formation. (Fig. 6A). It is interesting to note that a substantial proportion of Drosha migrated in the size range consistent with a single molecule of mCherry::DRSH-1 (150 kDa), even in the strain containing unaltered PASH-1::GFP, which may be indicative of a role for Drosha apart from the Microprocessor (Fig. 6A).

**Figure 6.**
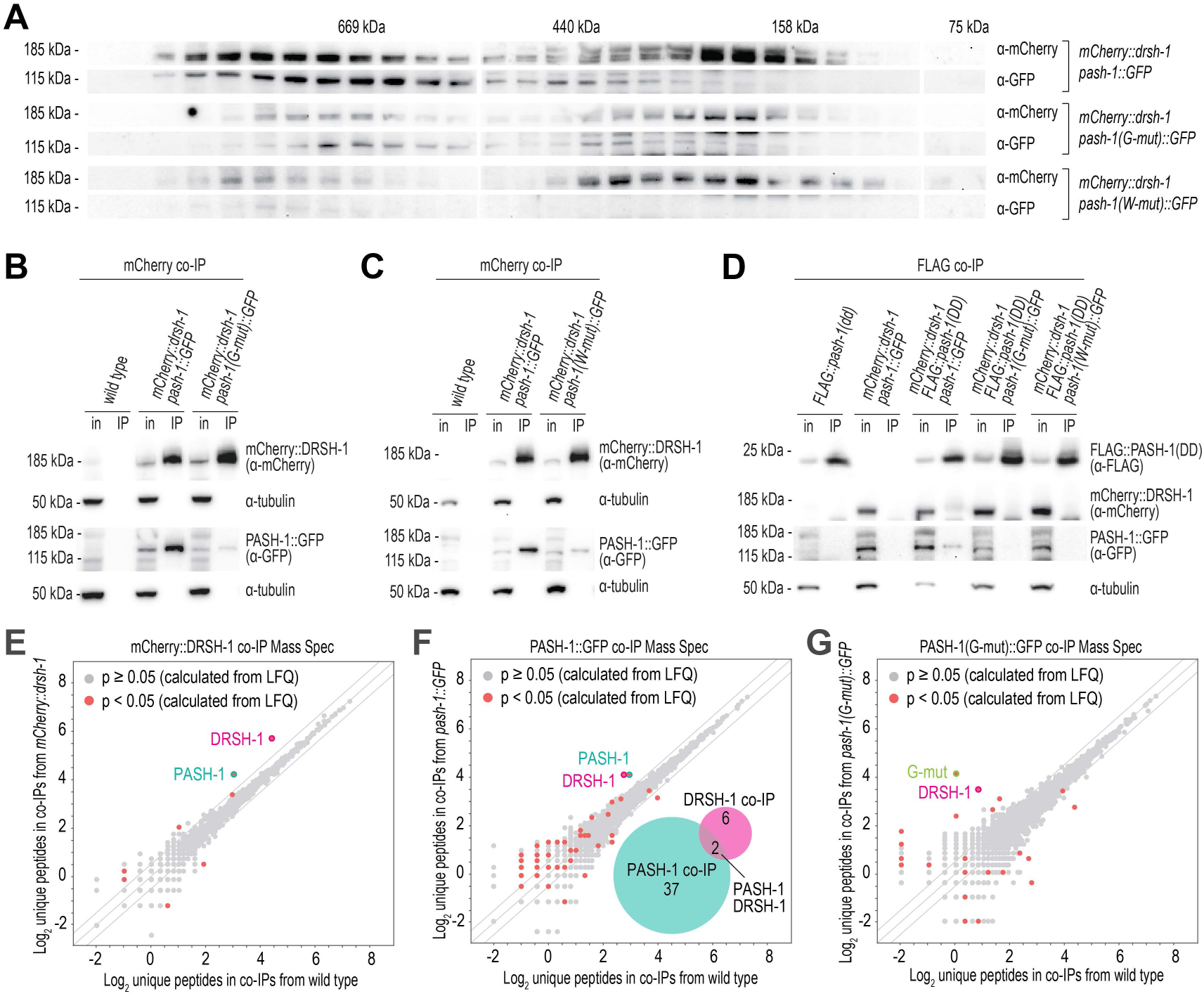
Microprocessor assembly is impaired in WW domain mutants. (A) Western blot analysis of unaltered and mutant PASH-1::GFP and unaltered mCherry::DRSH-1 from different protein fractions captured with size exclusion chromatography. Masses are approximated based on size markers. (B-C) Western blot analysis of PASH-1::GFP and PASH-1(G-mut)::GFP (B) or PASH-1 (W-mut)::GFP (C) co-lP’d with mCherry::DRSH-1. in, cell lysate input fraction; **IP,** co-IP fraction. Tubulin is shown as a loading control. Ratio IP to input was 9:1. (D) Western blot analysis of PASH-1 ::GFP, PASH-1(G-mut)::GFP, PASH-1(G-mut)::GFP, and mCherry::DRSH-1 co-lP’d with FLAG::PASH-1 (DD). Note that in these blots, 2.5x protein equivelents were loaded for the G-mut and W-mut co-lPs to make them directly comparable to unaltered PASH-1::GFP. Ratio IP to input was 20:1. (E-G) Mass spectrometry analysis of mCherry::DRSH-1 (E), PASH-1::GFP (F), and PASH-1(G-mut)::GFP (G) complexes from protein co-lPs. Scatter plots display the average 109_2_ unique peptide counts in co-lPs from wild type and the indicated transgenic strains. n = 4 biological replicates for each strain. Diagonal lines show 0-, 2-, and -2-fold enrichments. P values were calculated based on label free quantification (LFQ).

We then did reciprocal co-immunoprecipitation (co-IP) of Pasha and Drosha from animals containing wild type or the G or W mutant forms of Pasha fused to GFP, as well as wild type Drosha fused to mCherry. Non-mutant PASH-1::GFP co-IP’d with mCherry::DRSH-1 more efficiently than either the G or W mutants in each of three biological replicates (Fig. 6B-6C and Supplemental Fig. S6A-S6D). Similarly, mCherry::DRSH-1 co-IP’d more efficiently with non-mutant PASH-1::GFP than with either the G or W mutant forms (Supplemental Fig. S6A-S6D). Therefore, although not indispensable, the G and W residues contribute to the assembly or stability of the Microprocessor.

An expanded region of Pasha encompassing the WW domain is sufficient for dimerization *in vitro*^17^. It is therefore possible that the G and W residues promote Pasha dimerization, which could explain the loss of Microprocessor complex integrity in these mutants. To test this, we fused a 3xFLAG peptide to the PASH-1 dimerization region (DD) expressed under the control of ubiquitin (*ubl-1*) regulatory elements and integrated it into the *C. elegans* genome as a single-copy transgene, *FLAG::pash-1(DD)*^21^. We then introduced it by mating into strains containing *mCherry::drsh-1* and either *pash-1::GFP*, *pash-1(G-mut)::GFP,* or *pash-1(W-mut)::GFP* (Fig. 6D). Next, we co-IP’d FLAG::PASH-1(DD) and tested for its interaction with the unaltered and mutant forms of Pasha, as well as the unaltered form of Drosha. Unaltered PASH-1::GFP was readily detectable in FLAG::PASH-1(DD) co-IPs, demonstrating the dimerization domain identified *in vitro* also functions *in vivo* with full length Pasha (Fig. 6D and Supplemental Fig. S6E-S6F). In contrast, PASH-1(G-mut)::GFP and PASH-1(W-mut)::GFP were not detectable above background in FLAG co-IPs (Fig. 6D and Supplemental Fig. S6E-S6F). mCherry::DRSH-1 also co-IP’d with FLAG::PASH-1(DD) in the presence of non-mutant PASH-1::GFP, albeit at relatively low levels, but not with the G- or W-mutant forms, which did not associate with the DD fragment, indicating that one molecule of full-length Pasha is sufficient for association with Drosha (Fig. 6D and Supplemental Fig. S6E-S6F). These results demonstrate that the G and W residues are important for Pasha’s dimerization and proper assembly of the Microprocessor.

However, it is possible that the full heterotrimeric complex of one copy of Drosha and 2 copies of Pasha still forms in the mutants given that Drosha contains two binding sites for DGCR8/Pasha^42^. Regardless, the Pasha G and W mutants, with their reduced ability to dimerize, presumably form less stable and potentially less efficient Microprocessor complexes.

To identify the stable components of the Microprocessor and to determine if the composition changes in PASH-1(G-Mut)::GFP we did quantitative protein mass spectrometry. However, we did not identify any proteins that were enriched in both mCherry::DRSH-1 and PASH-1::GFP co-IPs aside from Drosha and Pasha using either label free quantification (LFQ; Supplemental Data 2) or unique peptide counts as a measurement (Fig. 6E-6F). This suggests that Drosha and Pasha are the only major stable components of the Microprocessor in *C. elegans*, although more transient interactors may also exist. Protein identified in in co-IPs from only one of the strains, mCherry::DRSH-1 or PASH-1::GFP, could be background and were not considered further. In Drosha and Pasha co-IPs from animals containing unaltered PASH-1::GFP we observed similar enrichment of unique peptides for both Drosha and Pasha (Fig. 6E-6F). In contrast, unique peptides of Pasha were enriched nearly 2-fold over unique Drosha peptides in co-IPs involving PASH-1(G-mut)::GFP, suggesting a weaker interaction between these proteins in the G-mutant, consistent with the Western blot analysis above (Fig. 6E-6G).

### Nuclear activity of the Microprocessor is essential for development

It is surprising that Pasha G mutants are relatively healthy and that W mutants are viable despite the loss of nuclear enrichment of the Microprocessor. This, together with our observation that unaltered PASH-1::GFP and mCherry::DRSH-1 are also expressed in the cytoplasm, raises the question of whether nuclear localization of the Microprocessor is essential for development in *C. elegans*. In humans, DGCR8/Pasha contains a nuclear localization signal (NLS) at its N terminus^34^. Although this region is poorly conserved in *C. elegans* and lacks an obvious NLS, we nevertheless deleted it in the *pash-1::GFP* line to determine if we could prevent Pasha’s nuclear localization and thereby assess whether nuclear activity of the Microprocessor is required for development (Fig. 7A). The deletion did result in a modest but variable reduction in nuclear enrichment, along with the occasional appearance of bright nuclear foci containing both mutant PASH-1(N-del)::GFP and mCherry::DRSH-1 (Fig. 7B and Supplemental Fig. S7). Animals containing the N-terminal deletion were relatively healthy, but like the G mutant were slightly dumpy and produced fewer viable progeny than non-mutant animals (Fig. 7C-7D). But because Pasha is still partially localized to the nucleus in these mutants, we could not conclude from these results if nuclear localization is essential for development.

**Figure 7.**
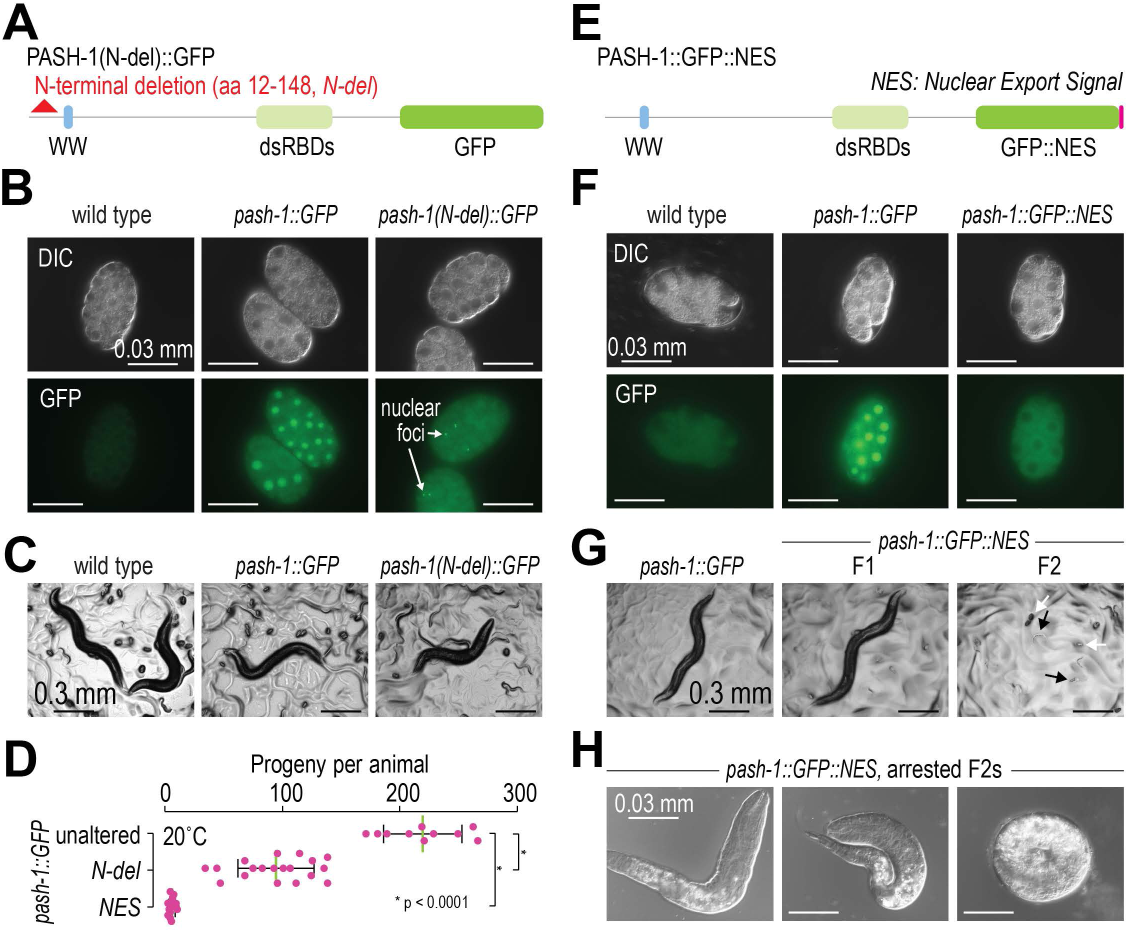
Nuclear localization of Pasha is required for development. **(A)** Diagram of the N-terminal PASH-1 deletion. (B) Representative images of PASH-1::GFP and PASH-1(N-del)::GFP expression in embryos. A wild type embryo lacking GFP is shown as a control. (C) Representative images of wild type, *pash-1 ::GFP,* and *pash-1(N-del)::GFP* animals. 1-2 gravid adults, larvae, and eggs are visible. (D) Numbers of progeny produced by wild type, *pash-1(N-del)::GFP,* and *pash-1::GFP::NES* animals grown at 20°C. n = **1**O (unaltered), n = 19 (N-del), and n = 14 (NES) animals. (E) Diagram of the PASH-1::GFP protein fused to a nuclear export signal (NES). (F) Representative images of PASH-1::GFP and PASH-1::GFP::NES expression in embryos. A wild type embryo lacking GFP is shown as a control. (G) Representative images of *pash-1 ::GFP* and *pash-1 ::GFP::NES* animals. **F1** animals are first generation segregates from a heterozygous parent. F2 are descended from a homozygous parent. White arrows point to arrested embryos and black arrows point to arrested larvae. (H) Arrested *pash-1 ::GFP::NES* larvae and embryos.

We then introduced a nuclear export signal (NES) sequence at the 3’ end of *pash-1::GFP* (*pash-1::GFP::NES;* Fig. 7E)^43,44^. The NES caused strong mislocalization of PASH-1::GFP to the cytoplasm, thereby allowing us to assess the requirement for nuclear localization (Fig. 7F). *pash-1::GFP::NES* mutants displayed a high incidence of embryonic arrest and only occasionally hatched into larvae, which also arrested early in development, although like other *pash-1* mutant alleles, the homozygous mutant F1 progeny of heterozygous animals were healthy (Fig. 7D and 7G-7H). These results underscore the critical requirement for nuclear processing of pri-miRNAs and thus the importance of nuclear localization of the Microprocessor.

## DISCUSSION

*C. elegans* has pioneered miRNA exploration since the discovery of these tiny RNAs in this genetically tractable whole animal system^45,46^. The pri-miRNA sensor developed here leverages the awesome genetic malleability and translucent appearance of *C. elegans* to enable whole animal genome-wide screens exploring the first step in miRNA biogenesis. In a proof-of-principal study, we identified numerous independent lines in which the sensor was desilenced, focusing on the one containing the Pasha G>R mutation described here. Undoubtedly, further genetic analysis with the sensor will uncover additional features of primary miRNA processing. Thus, we believe that the sensor will be a valuable resource for exploring the miRNA pathway. Furthermore, because the sensor is exceptionally easy to screen using a simple stereo microscope with fluorescence capabilities, it can be easily incorporated into both research and teaching laboratories. The sensor strain is available from the Caenorhabditis Genetics Center.

Our initial screen to identify cis-acting mutations affecting pri-miRNA processing (i.e. mutations in the pir-miRNA sensor) was less productive than expected, with only 1 in 500 candidates having a mutation within the miRNA hairpin region. It is possible that our screen lacked sufficient depth or that pri-miR-58 processing is resistant to single point mutations. Unlike mammals and at least some insects, *C. elegans* does not have the obvious sequence motifs that promote pri-miRNA recognition and processing, which might otherwise emerge from forward genetic screens^15,47–49^. Lack of such sequence elements could be compensated for in part by alternative dsRNA recognition sites^50^. In flies, pri-miRNAs lacking optimal sequence motifs depend on an interaction between Pasha and RNA Pol II that may aid in their recognition by the Microprocessor^25^. While it is possible that this mechanism is also important for miRNA processing in worms, we did not detect an interaction between Pasha and Pol II. Perhaps miRNA hairpin structure is adequate for efficient recognition by the Microprocessor in worms. Consistent with this possibility the one cis-acting mutation we identified in the hairpin (changing a G-C base pair to G-U), leads to what is likely a very modest structural change in the hairpin. Because the mutation occurs within the miRNA region and this position is not enriched for a C nucleotide in *C. elegans* miRNAs it is unlikely that a C is itself important. In humans, mismatches in the upper stem of the miRNA hairpin, near the region identified here, can also impact Drosha cleavage^51,52^. Such mismatches can also function positively to promote cleavage at the correct position^51,53^. However, because the pri-miR-58 sensor does not report on processing accuracy, the mutation identified here must lead to a reduction in cleavage efficiency, which could also be accompanied by a reduction in accuracy, which we did not explore. Thus, consistent with results in humans, the miRNA duplex region is important for miRNA recognition or processing.

We uncovered a crossregulatory interaction between Pasha and Drosha important for their localization to the nucleus. This interaction is facilitated by dimerization of Pasha via the GW motif of its WW domain. Our findings suggest that entry or retention of Pasha or Drosha in the nucleus is dependent on association with an intact Microprocessor complex. This was unexpected because both proteins appear to have their own nuclear localization signals, at least in mammals^34,35^. However, the N terminal regions of both DGCR8/Pasha and Drosha are poorly conserved in *C. elegans* but nonetheless the loss of nuclear enrichment when we deleted the N terminus of Pasha suggests that it may have a cryptic NLS. It is possible that in *C. elegans*, Pasha and Drosha both have weak NLSs and their combined function is important for the nuclear localization of the complex. Alternatively, protein modifications may be important for nuclear localization and such modification could depend on an intact complex. In humans, for example, phosphorylation of Drosha promotes its nuclear localization^35^. Regulation of nuclear localization could provide a valuable control mechanism to prevent Microprocessor-independent functions for Pasha or Drosha from occurring in the nucleus. Pasha has been shown to regulate neuronal genes in the absence of Drosha in *Drosophila* so there is certainly a precedence for the individual Microprocessor components to function independently of each other, which could have unintended consequences if not tightly regulated^54^. Furthermore, our protein fractionation experiments suggest that a large fraction of Drosha is not in complex with Pasha and Pasha in turn may also exist in a complex that doesn’t include Drosha. It is interesting to note here that we did not identify a clear post-transcriptional gene regulatory mechanism between Pasha and Drosha like that of mammals^20^. However, it is possible that crossregulation of Pasha and Drosha nuclear localization serves a similar role in *C. elegans*.

Despite a crucial role for Pasha in *C. elegans* development, we found that mutations within Pasha’s WW domain are surprisingly well tolerated. Interestingly, the W mutant is sick, whereas the G mutant is relatively healthy despite the two residues having a similar requirement for Microprocessor assembly and nuclear localization. This suggests that defects in assembly and nuclear localization of the Microprocessor only partially explain the phenotype of W mutants. Presumably the W residue has a more critical role in miRNA processing than the G residue, but whether this reflects a distinct role is unclear. The W residue may have additional roles, such as interactions with other proteins important for miRNA processing. Presumably these interactions are weak or transient as we did not identify them by mass spectrometry of Pasha interactors. Alternatively, the W residue could have a more critical role in Pasha’s association with, or orientation on, pri-miRNAs.

It is well-established that pri-miRNA processing occurs in the nucleus and that Drosha and DGCR8/Pasha both localize to the nucleus^8,33,36^. Pri-miRNA processing is thought to occur co-transcriptionally, facilitated in part by Drosha’s N-terminus, which contains a proline-rich disordered domain^55–57^. Interestingly, the first ∼220 amino acids of Drosha’s N terminus, containing most of the proline-rich region, is completely absent in *C. elegans*^57^. Furthermore, while most miRNAs occur in the introns of protein-coding or non-coding genes in humans, only a small subset are produced from intronic regions in *C. elegans*. Even the intronic miRNAs that do exist in *C. elegans* appear to have regulatory elements distinct from the host transcript^58^. Therefore, it is possible that, unlike in mammals, *C. elegans* miRNAs are predominantly processed post-transcriptionally. Nevertheless, our results showing that exclusion of Pasha from the nucleus leads to embryonic or early larval lethality suggests that pri-miRNA processing, regardless of whether it is co-transcriptional or post-transcriptional, must occur in the nucleus. It is still possible that some level of pri-miRNA processing occurs in the cytoplasm, but if so, it is unlikely that it is sufficient to rescue loss of nuclear processing. However, it is also possible that the Microprocessor serves other functions that underlie the lethality that occurs in its absence from the nucleus.

It is interesting to note that all the Pasha mutants generated in this study that develop into adults display a mildly squatty or dumpy phenotype, suggesting that the underlying regulatory network is highly sensitive to a modest loss of miRNAs. This contrasts with other processes controlled by miRNAs, such as developmental timing, embryogenesis, vulval development, and fertility which are robust to the modest loss of miRNAs observed in Pasha G and N-del mutants. To our knowledge, the dumpy phenotype, although relatively well studied in *C. elegans*, has not been linked to loss of miRNA activity. It is possible that mislocalization of the Microprocessor to the cytoplasm leads to mis-silencing of genes involved in controlling body shape, such as cuticular genes. While it is probable that this relates to miRNAs either directly or indirectly, it could instead be attributable to a non-miRNA role of Pasha or the Microprocessor.

The model that emerges from this study suggests that Microprocessor integrity, partially facilitated by the dimerization of Pasha through the GW motif in the WW domain, is important for nuclear entry or retention, which in turn is necessary for miRNA processing and proper development. It will be important in future studies to determine if the WW domain is important for interaction with other proteins in *C. elegans* using more sensitive techniques then those applied here. It also remains to be seen whether miRNA processing in *C. elegans* is co-transcriptional or if it occurs post-transcriptionally, differently from the mechanism in mammals. Additionally, it will be interesting in future studies to examine potential cytoplasmic functions of the Microprocessor and roles for Pasha and Drosha that do not depend on the Microprocessor. Genetic studies, such as those using the pri-miR-58 sensor developed in this research, are expected to reveal new factors involved in miRNA processing and additional roles for Pasha and Drosha, enhancing our understanding of these critical processes.

## METHODS

### Strains

To generate the pri-miR-58 sensor, *ram3[(pCMP1)ubl-1::mCherry::pri-mir-58::ubl-1 + Cbr-unc-119(+)]* II, 3000 bp upstream of the *ubl-1* start codon was amplified from wild type *C. elegans* (N2) genomic DNA using Phusion polymerase (ThermoFisher, F534L) and cloned into pDONR 221 P1-P4 (Addgene plasmid #186351) using Gateway Recombination Cloning Technology (ThermoFisher), as described^59^. mCherry was PCR amplified from plasmid DNA with the primers GGGGACAACTTTTCTATACAAAGTTGACATGGTCTCAAAGGGTGAAGAAG and GGGGACAACTTTATTATACAAAGTTGTTTACTTATACAATTCATCCATG and cloned into pDONR 221 P4r-P3r (Addgene plasmid #121527). The miR-58 hairpin sequence and 100 bp upstream and 1000 bp downstream was amplified from wild type genomic DNA and cloned into pDONR 221 P3-P2. The entry plasmids were recombined into the destination vector CMP1 and then transformed into *C. elegans* strain EG6699 using Mos1-mediated single copy insertion as described^21,59^. The *ubl-1::mCherry::pri-mir-58-sensor-mut* mutation was introduced into the CMP1 *ubl-1::mCherry::pri-mir-58-sensor* plasmid using NEB Q5 site directed mutagenesis (New England Biolabs, E0554S) with the primers GGGATGAGATTGTTCAGTACG and TATGGTATTGGACGAAGTG. *FLAG::pash-1(DD)* was PCR amplified from cDNA with primers extending the 5’ end with a 3xFLAG peptide sequence and the 5’ and 3’ ends with pDONR entry vector compatible sequences (GGGGACAACTTTTCTATACAAAGTTGACATGGATTATAAAGACGATGACGATAAGCGTGACTACAAGGACGACGACGACAAGCGTGATTACAAGGATGACGATGACAAGAGTGTCGGTGAACAAATTCG and GGGGACAACTTTATTATACAAAGTTGTCGGTGACATTCGAAGCTCCTTGAC). The resulting fragment was cloned into pDONR 221 P4r-P3r and then recombined into CMP1 with the *ubl-1* regulatory sequences described above. A stop codon within the *ubl-1* 3’UTR was utilized to terminate translation, extending the C terminus of 3xFLAG::PASH-1dd protein by 22 amino acids. The resulting plasmid was transformed into wild-type (N2) worms using Mos1-mediated single copy insertion as described^21,59^. VC1138[*drsh-1(ok369) I/hT2[bli-4(e937) let-?(q782) qIs48*](I;III)], MT15024[*mir-58(n4640)* IV], and KW2090[*cdk-9(tm2884) I/hT2 [bli-4(e937) let-?(q782) qls48]* (I;III)] were previously described^22,38,60^. TAM93[*pash-1(ram33)* I] containing the 38a mutation that was identified in a forward genetic screen was outcrossed three times to wild type (N2). PHX4329[*pash-1(syb4327)* I], PHX6091[*pash-1(syb6091)* I], PHX6157[*pash-1(ram33 syb6157)* I], PHX6655[*pash-1(syb6091 syb6655)* I], PHX6628[*drsh-1(syb6628)* I], PHX7689[*pash-1(syb66091 syb7689)* I], PHX8549[*pash-1(syb6091 syb8549)* I], PHX8348[*pash-1(syb8348)/hT2[bli-4(e937) let-?(q782) qIs48]* (I;III)], and PHX6604[*pash-1(syb6091 syb6604)* I] (miR-71 seed target edited from TCTTTCA to AGAAAGT) were generated by Suny Biotechnology (Fuzhou, China) using CRISPR-Cas9 genome editing and outcrossed to wild type (N2) or to each other to generate combinatorial strains (Supplemental Data 3)^26–29^.

### Animal growth conditions

For the mass spectrometry experiment, animals were expanded on egg plates for one generation and then transferred to NGM plates as described^61^. For all other experiments, animals were grown on NGM plates containing *E. coli* OP50 at 20°C unless noted otherwise.

### Forward genetics

L4 and young adult *C. elegans* were rotated in M9 buffer solution containing 1% ethyl methyl sulfonate (EMS) solution for 4 hours at room temperature and then washed 4x in M9. 50 animals were plated onto 10 cm NGM plates containing *E. coli* OP50 and their F1 (cis-factor screen) or F2 (general screen) progeny were selected for bright red fluorescence. For the cis-factor screen, individual bright fluorescent animals were lysed and the miR-58 hairpin sequence of the *pri-miR-58* sensor was sequenced using Sanger sequencing. For the general screen, individual bright fluorescent animals were selected and expanded for several generations. The brightest healthy line after several generations was subjected to whole genome sequencing to identify the causal mutation.

### Genome sequencing

DNA from EMS line 38a was isolated using the Gentra Puregene Tissue Kit (QIAGEN, 158667). Libraries were prepared and sequenced using Novogene’s Whole Genome Sequencing service, which utilizes the NEBNext DNA Library Prep Kit, and sequenced on an Illumina NovaSeq (PE150). Data was processed and single nucleotide polymorphisms (SNPs) were identified using Novogene’s Bioinformatic analysis pipeline. Briefly, reads were aligned to the *C. elegans* genome (WS190) using BWA (parameters: ’mem -t 4 -k 32 -M’)^62^. PCR duplicates were removed using SAMTOOLS^63^. SNPs were called using GATK (parameters: ’-T HaplotypeCaller --gcpHMM 10 -stand_emit_conf 10 - stand_call_conf 30’)^64^. Results were filtered to reduce the error rate (parameters: ’QD < 2.0 || FS > 60.0 || MQ < 30.0 || HaplotypeScore > 13.0 || MappingQualityRankSum < -12.5 || eadPosRankSum < -8.0’). SNPs were annotated using ANNOVAR^65^.

### RNA isolation

Gravid adult animals, grown for 54 hours at 25°C or 72 hours at 20°C after L1 synchronization, were washed off plates and then washed 3x in M9 buffer and flash frozen in liquid nitrogen. RNA was isolated from whole gravid adult animals using Trizol (Life Technologies, cat# 15596018) according to the manufacturer’s recommendations but with a second chloroform extraction step. For mRNA qRT-PCR, total RNA was DNase-treated using the TURBO DNA-free Kit (ThermoFisher, AM1907) according to the manufacturer’s recommendations.

### qRT-PCR

Small RNA quantitative real-time-PCR was done with TaqMan reagents and custom probes targeting miR-1 (Life Technologies, assay name: hsa-miR-1, 4427975), miR-58a-3p (sequence: TGAGATCGTTCAGTACGGCAAT), miR-35-3p (sequence: TCACCGGGTGGAAACTAGCAGT), 21UR-1 (sequence: TGGTACGTACGTTAACCGTGC), and 22G-rRNA (sequence: GAAGAAAACTCTAGCTCGGTCT) following the manufacturer’s recommendations (Life Technologies, 4331348). Pri-miR-58 qRT-PCR was done with custom TaqMan probe sets targeting *pri-miR-58* and *act-1*. *pash-1* and *drsh-1* mRNA qRT-PCR analysis was done using SYBR Green and the following primers: *pash-1*, GTTCACTCGTGTCGTCACTC and CGTTTTCGTGCAGCTCATCC; *drsh-1*, GTACTTGGAATCGAAGGACC and AGATTAGCCAAAGCCAGCTC; *rpl-32* (housekeeping gene), CATGAGTCCGACAGATACCG and ACGAAGCGGGTTCTTCTGTC. Ct values were captured using a CF96 Real-Time PCR Detection System (Bio-Rad) and averaged across three technical replicates for each of 3 biological replicates. The 2-ddCt method was used to calculate relative fold changes between conditions^66^. Two-sample t-tests were used to calculate P values.

### Small RNA sequencing and data analysis

Small RNA sequencing libraries were prepared using the NEBNext Multiplex Small RNA Library Prep Set for Illumina following the manufacturer’s recommendations with exclusion of the initial size selection and the 3’ ligation step changed to 16°C for 18 hours to improve capture of methylated small RNAs (New England Biolabs, E7300S). Small RNA PCR products were size selected on a 10% polyacrylamide gel, transferred by electrophoresis to DE81 chromatography paper, eluted at 70°C for 20 minutes in the presence of 1 M NaCl, and precipitated at -80°C overnight in the presence of 13 ug/ml glycogen and 67% EtOH. End-to-end data analysis was done using the default configuration in tinyRNA with the *C. elegans* genome WS279 release^67,68^.

### Microscopy

For imaging PASH-1::GFP and mCherry::DRSH-1, adult whole animals and embryos were mounted on glass slides containing 1.5% Agarose pads and imaged on an Axio Imager Z2 Microscope (Zeiss) after immobilization in 25 uM sodium azide. mCherry fluorescence signal quantification was done in Fiji^69^. For imaging *pri-mir-58* sensor-transgenic animals, a Discovery V8 Stereo Microscope (Zeiss) was used.

### RNAi

For RNAi knockdown, synchronized L1 larvae were placed on RNAi plates containing IPTG and *E. coli* HT115 expressing dsRNA matching *cdk-9*, *ama-1*, *alg-1/alg-2, pash-1*, *drsh-1*, or empty vector (L4440) and were grown at 20°C^70,71^. Where applicable, gravid adults were collected for protein and RNA isolation after 72 hours of treatment.

### Protein co-immunoprecipitation

PASH-1::GFP, PASH-1(G-mut)::GFP, PASH-1(W-mut)::GFP, FLAG::PASH-1(DD), mCherry::DRSH-1, and RNA Polymerase II were co-IP’d from three biological replicate samples with ∼12,000 gravid adult animals each. Animals were washed from plates and then 3x in M9 salt buffer, frozen in liquid nitrogen, and lysed in 1.2 ml 50 mM Tris-Cl, pH 7.4, 100 mM KCl, 2.5 mM MgCl2, 0.1% Igepal CA-630, 0.5 mM PMSF, and 1X Pierce Protease Inhibitor Tablets (Pierce Biotechnology, cat# 88266). Cell lysates were cleared for 10 min at 12,000 RCF at 4°C. Cleared lysates were split into input and co-IP fractions. For GFP and mCherry co-IPs, cell lysates were incubated with 25 ul ChromoTek GFP-Trap Magnetic Agarose Beads (Proteintech, gtma-100) (GFP co-IPs) or ChromoTek RFP-Trap Magnetic Agarose Beads (Proteintech, rtma-20) (mCherry co-IPs) for 1 hour at 4°C. For FLAG co-IPs, lysates were incubated with 7 ug FLAG antibody (Sigma, F3165) for 1 hour at 4°C. After 30 minutes of FLAG antibody incubation, 60 ul of Protein A agarose beads (Roche, PROTAA-RO) were added. For Pol II-PASH-1::GFP co-IPs, Pol II was co-IP’d with 4 ug Pol II antibody (Santa Cruz Biotechnology, sc-537117) and PASH-1::GFP was co-IP’d with 5 ug GFP antibody (Invitrogen, A11120) for 90 minutes at 4°C. After 45 of GFP antibody incubation, 50 ul of SureBeads Protein G magnetic agarose beads (Bio-Rad, 1614023) were added. Beads were washed three times in lysis buffer and protein was eluted at 95°C for 5 minutes in 1X SDS Blue Loading Buffer containing 50 mM DTT.

### Size exclusion chromatography

Animals were prepared as described above for co-IPs. Cell lysates were treated with 0.5 ug/ul RNase I for 1 hour at 4°C. Proteins were separated on a Superdex 200 Increase small-scale SEC columns, 10/300 GL (Cytiva, 28990944) according to the manufacturer’s recommendations. Western blot analysis of the fractions was done as described below.

### Western blots

Co-IP’d proteins, cell lysates, and SEC fractions were resolved on Bolt 15-well 4-12% Bis-Tris Plus Gels (Life Technologies, NW04125BOX), transferred to nitrocellulose membranes, and probed with GFP (Santa Cruz Biotechnology, sc-9996 HRP), mCherry (Proteintech, 6G6), FLAG (Pierce, PA1-984B), Pol II (Santa Cruz Biotechnology, sc-537117), tubulin (Abcam, ab40742), and actin (Abcam, ab3280) antibodies. Blots were imaged and quantified on a FluorChem E Imaging System (ProteinSimple).

### Mass spectrometry

Gravid adult animals (200 ul packed worms per replicate, 4 biological replicates per sample) were washed from NGM plates and then 3x in M9. Animals were flash frozen in liquid nitrogen, thawed, and lysed by sonication (10 cycles of 30 sec ON, 30 sec OFF, high efficiency) in lysis buffer (25 mM Tris HCl pH 7.5, 150 mM NaCl, 1.5 mM MgCl2, 1 mM DTT, 0.1% Triton X-100, and 1 tablet/40 ml cOmplete Mini, EDTA-free protease Inhibitor cocktail). Cell lysates were cleared for 10 min at 21,000 RCF at 4°C. BCA assays were used to normalize the starting protein amount. 20 ul ChromoTek GFP-Trap Magnetic Agarose Beads (Proteintech, gtma-100) or ChromoTek RFP-Trap Magnetic Agarose Beads (Proteintech, rtma-20) were washed 3x in 500 ul wash buffer (25mM Tris HCL pH 7.5, 150 mM NaCl, 1.5 mM MgCl2, and 1 mM DTT) and then incubated with cleared cell lysates overnight while rotating at 4°C. Beads were washed 3x in wash buffer and resuspended in 50 ul in 1X LDS Sample Buffer and incubated at 95°C for 10 min to elute protein. Mass spectrometry and data analysis were done as described^61^.

### Fertility assays

Individual animals were grown from L1 stage larvae on OP50 at 20°C or 25°C and their progeny were summed every day until the cessation of egg laying. Progeny counts included all larval stage animals.

### Statistics

DESeq2 was used within the tinyRNA sRNA-seq data analysis workflow to perform statistical analysis using the Wald test^67,72^. R and GraphPad Prism were used to do ANOVA and TukeyHSD or Dunnett tests for fertility assays. Microsoft Excel and GraphPad Prism were used to perform t-tests for qRT-PCR data analysis and in comparing total iso-miR and pre-miRNA read levels.

### Data availability

Raw and processed sequencing data generated in this study has been submitted to the NCBI Gene Expression Omnibus (GEO; https://www.ncbi.nlm.nih.gov/geo/) under accession number GSE263914.

## ACKNOWLEDGEMENTS

Jiaxuan Chen is gratefully acknowledged for his help with mass spectrometry and data analysis (instrumentation funded by the German Research Foundation, DFG INST 247/766-1 FUGG). Thanks also to Alivia Ball and Maritza Soto-Ojeda for help with media and solutions. Strains not generated in this study were provided by the Caenorhabditis Genetics Center (CGC), which is funded by the National Institutes of Health Office of Research Infrastructure Programs (P40 OD010440). The VC1138 strain was generated by the C. elegans Reverse Genetics Core Facility at the University of British Columbia, which is part of the international C. elegans Gene Knockout Consortium. This work was supported by the National Institutes of Health [R35GM119775 to T.A.M.], the Institute of Molecular Biology - Meinz [core funding to R.F.K.], and the German Research Foundation [DFG grant 252386272 to R.F.K.].

## Supplemental Figures

**Supplemental Figure S1.**
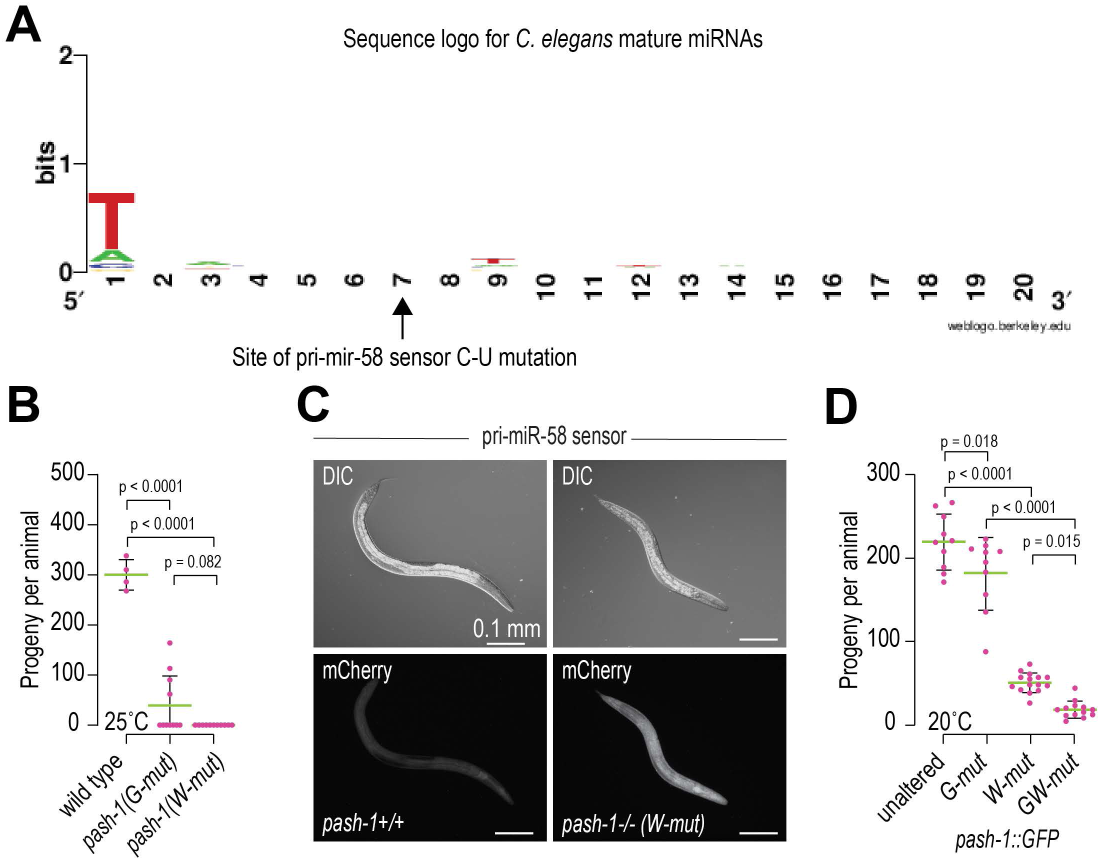
Fertility and pri-miRNA processing defects in Pasha WW domain mutants. (A) The Weblogo shows conservation across the first 20 nucleotides of C. elegans miRNAs (miRbase release 22). (B) Numbers of progeny produced by wild type and *pash-1(ram33)* (G-mut) and *pash-1(syb4327) (W-muO* animals grown at 25°C. Error bars are standard deviation from the mean of 4 (wild type) or 11 animals. (C) Representative images showing mCherry fluorescence in control (non-mutated) or *pash-1(syb4327) (W-mut)* animals containing the pri-miR-58 sensor construct. DIC and mCherry fluorescence images are shown. (D) Numbers of progeny produced by *pash-1::GFP, pash-1(G-mut)::GFP, pash-1(W-mut)::GFP,* and *pash-1(GW-mut)::GFP* animals grown at 20°C. Error bars are standard deviation from the mean of 10-14 animals.

**Supplemental Figure S2.**
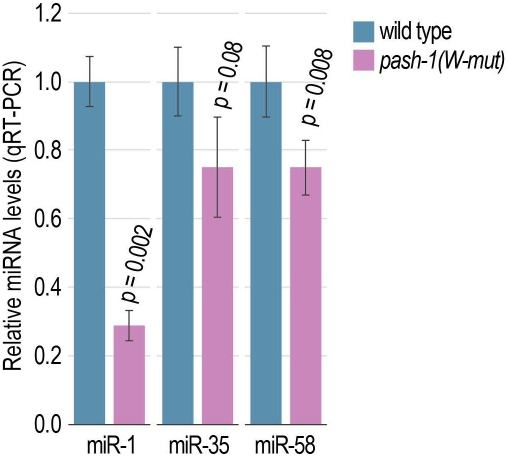
Reduction in miRNA levels in Pasha W mutants. Relative levels of miR-1, miR-35, and miR-58 in wild type and *pash-1(syb4327) (W-mut)* grown at 20°C as determined TaqMan qRT-PCR. Data is normalized to 21UR-1 piRNA levels. Error bars are standard deviation from the mean of 3 biological replicates. P values are for comparison to wild type.

**Supplemental Figure S3.**
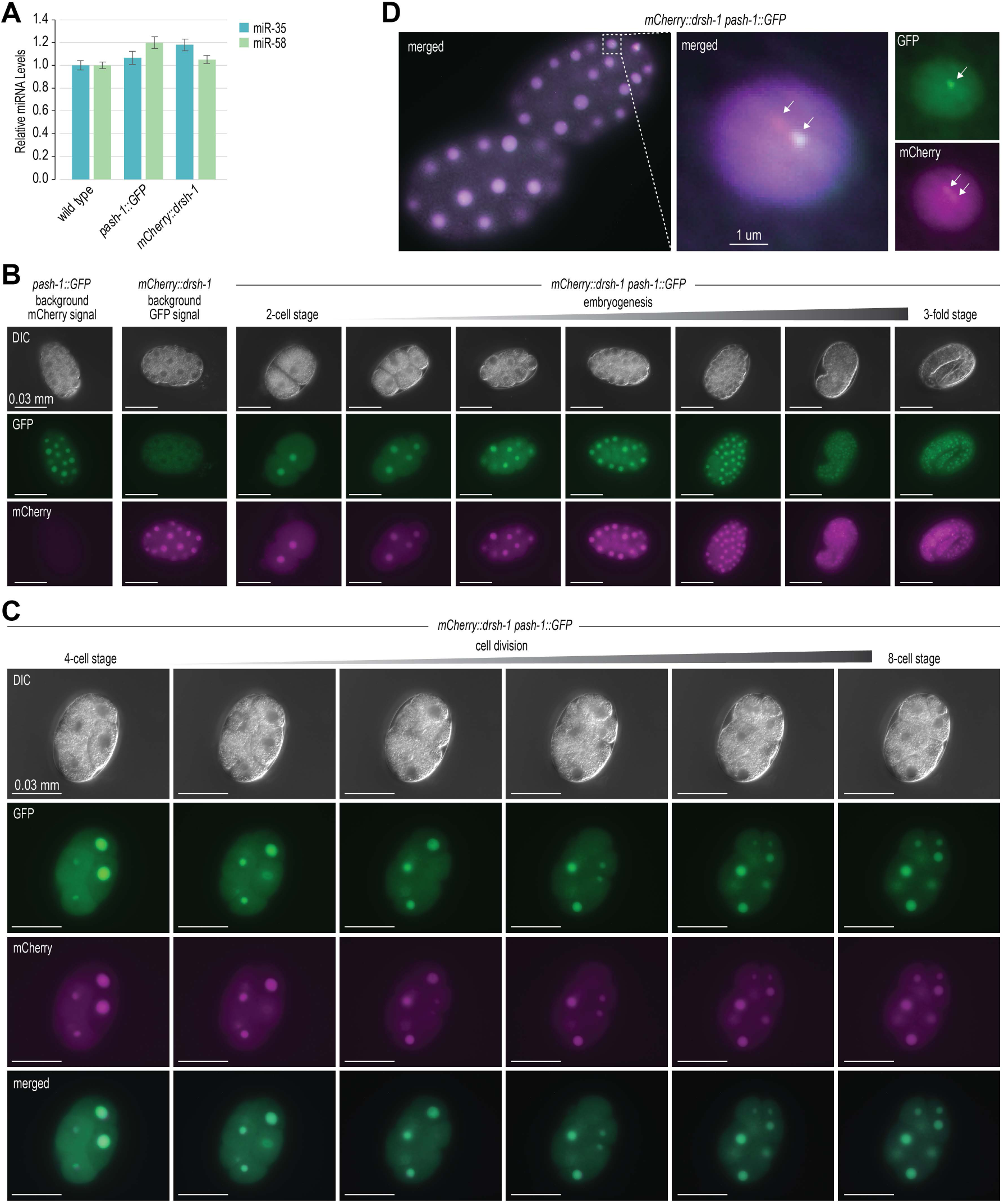
Subcellular localization of Pasha and Drosha. (A) Relative levels of miR-35 and miR-58 in wild type, *pash-1 ::GFP,* and *mCherry::drsh-1* grown at 20°C as determined TaqMan qRT-PCR. Data is normalized to 21UR-1 piRNA levels. (B) Representative images of PASH-1::GFP and mCherry::DRSH-1 expression in 2-cell through 3-fold stage embryos. (C) Representative images of PASH-1 ::GFP and mCherry::DRSH-1 localization during cell divisions taking place between 4-cell and 8-cell stage embryos. DIC, GFP, mCherry, and GFP-mCherry merged images are shown. (D) PASH-1 ::GFP and mCherry::DRSH-1 nuclear localization in a single nucleus from an embryo. Arrows point to nuclear foci.

**Supplemental Figure S4.**
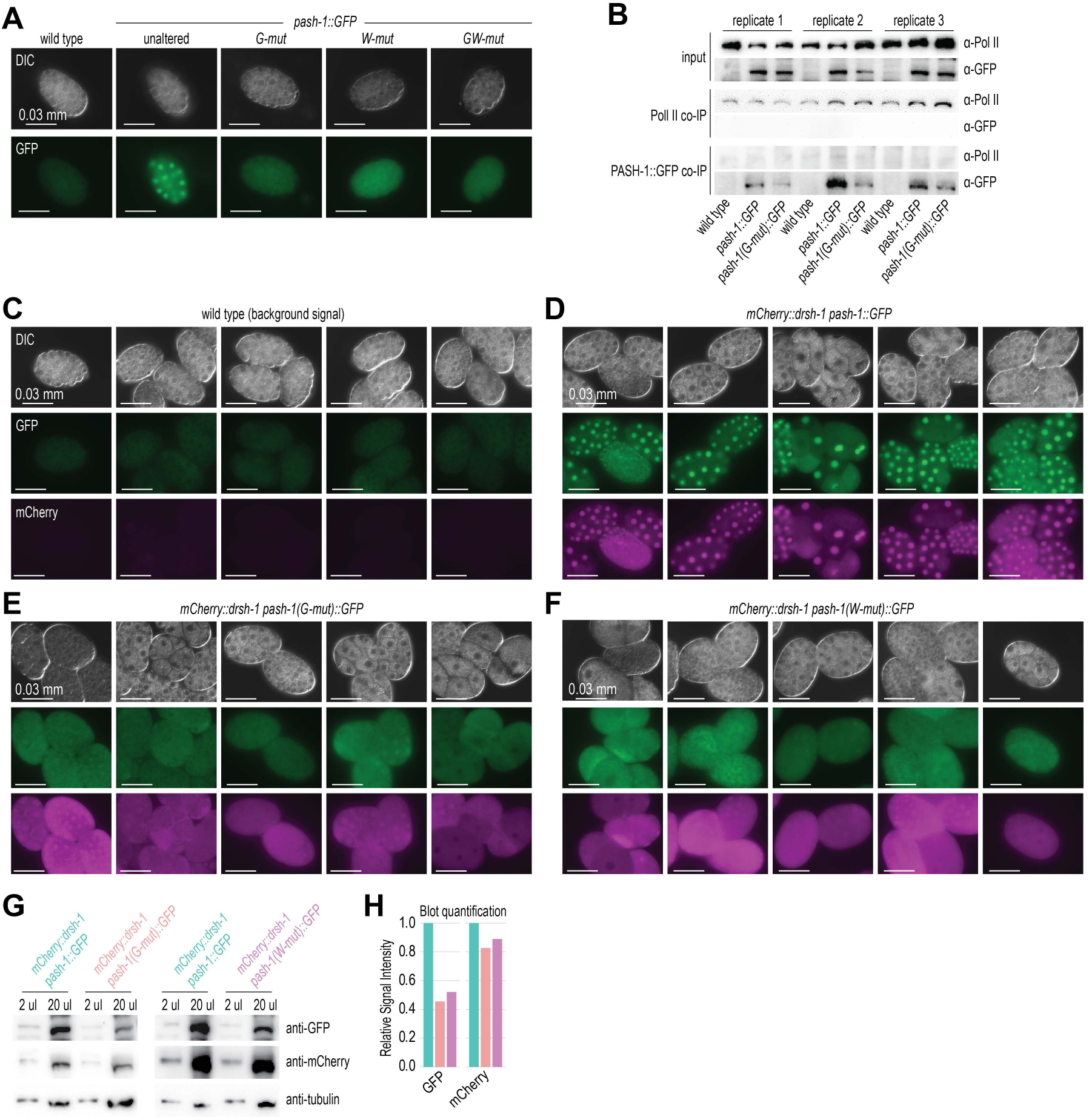
Loss of nuclear enrichment of the Microprocessor in Pasha WW domain mutants. (A) Representative images of wild type and *pash-1 ::GFP, pash-1(G-mut)::GFP, pash-1 (W-mut)::GFP,* and *pash-1(GW-mut)::GFP* embryos. (B) Western blot analysis of PASH-1 ::GFP and Pol II co-lPs. 3 biological replicates are shown. (C-F) Images of wild type embryos (B) and *mCherry::drsh-1 pash-1::GFP* (D), *mCherry::drsh-1 pash-1(G-mut)::GFP* (E), *mCherry::drsh-1 pash-1(W-mut)::GFP* (F) transgenic embryos. DIC, GFP, and mCherry images are shown. (G) mCherry::DRSH-1, PASH-1::GFP, PASH-1(G-mut)::GFP, and PASH-1(W-mut)::GFP protein levels as determined by western blot. Tubulin is shown as a loading control. (H) Quantification of the blot in (G).

**Supplemental Figure S5.**
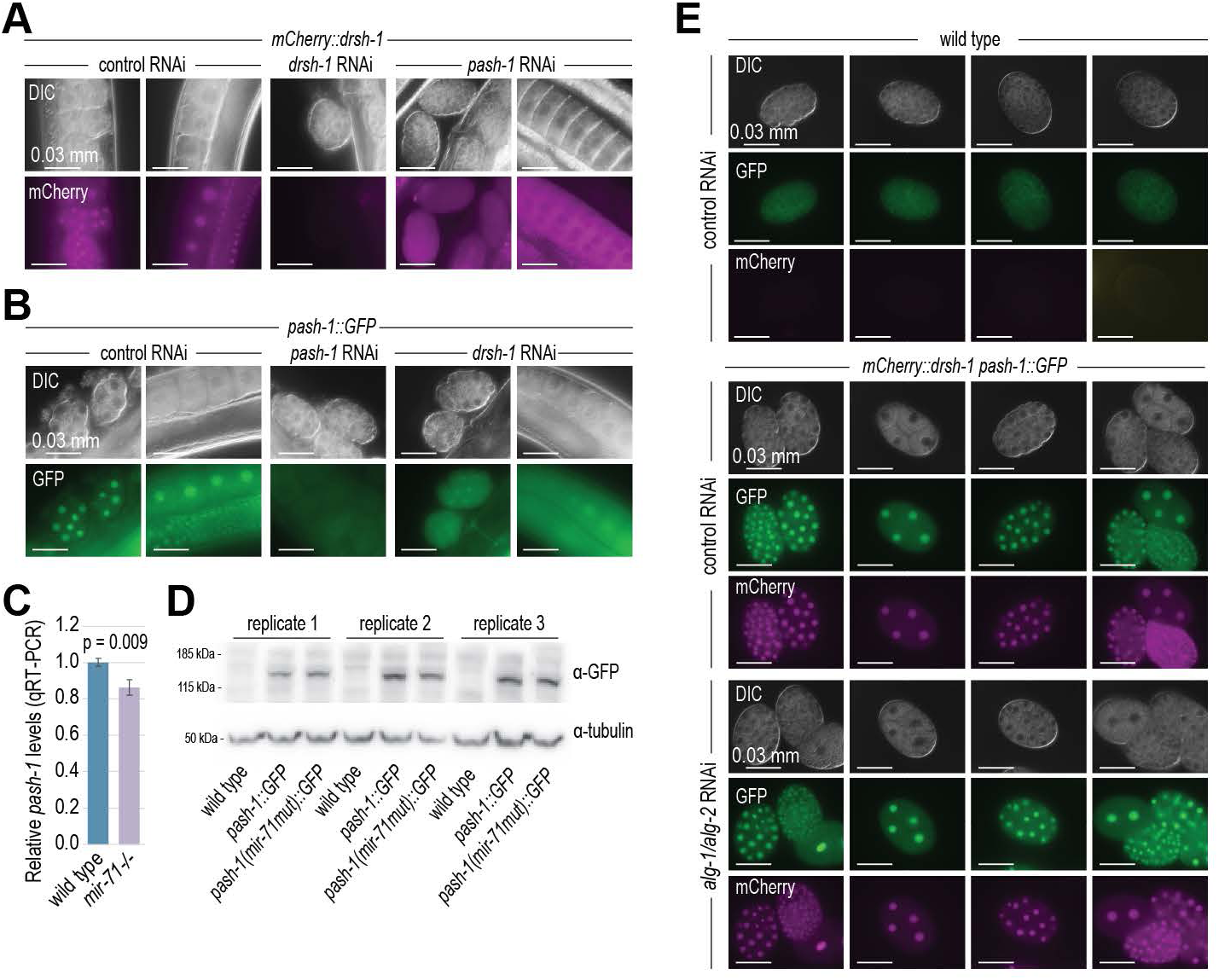
Pasha and Drosha require each other for proper nuclear localization. (A) Representative images showing mCherry::DRSH-1 localization in germlines and embryos of control (L4440), *drsh-1,* and *pash-1* RNAi-treated animals. DIC and mCherry images are shown. (B) Representative images showing PASH-1 ::GFP localization in germlines and embryos of control (L4440), *drsh-1,* and *pash-1* RNAi-treated animals. DIC and GFP images are shown. (C) Relative *pash-1* mRNA levels in wild type and *mir-71-1-* gravid adult animals, as determined by qRT-PCR. *rpl-32* mRNA levels were used for normalization. Error bars are standard deviation from the mean for 3 biological replicates. (D) PASH-1 protein levels produced from *pash-1 ::GFP* and *pash-1(mir-71mut)::GFP.* 3 biological replicates are shown in the Western blot. Tubulin was used as a loading control. (E) Representative images showing mCherry::DRSH-1 and PASH-1 ::GFP expression in embryos of control (L4440) and *alg-1/alg-2* RNAi-treated animals. DIC, GFP, and mCherry images are shown.

**Supplemental Figure S6.**
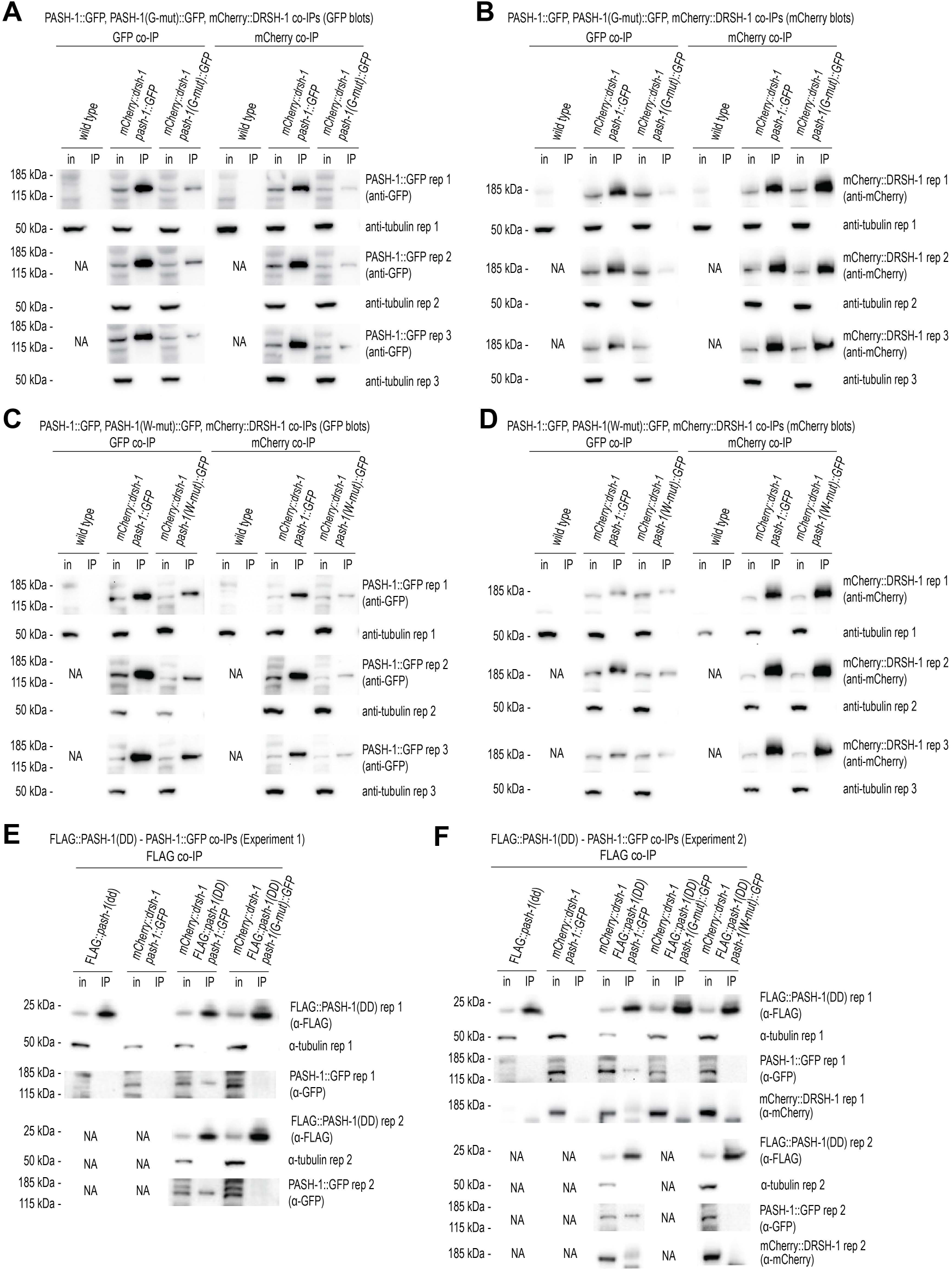
Pasha and Drosha interaction in Pasha WW domain mutants. (A-D) Western blot analysis of Pasha-Drosha interactions. Ratio of IP to input was 9:1. (A) Western blot analysis of PASH-1::GFP and PASH-1(G-mut)::GFP co-lP’d with mCherry antibody (recognizing mCherry::DRSH-1) or GFP antibody. in, cell lysate input fraction; IP, co-IP fraction. Tubulin is shown as a loading control. 3 biological replicates are shown, except for wild type. (B) Western blot analysis of mCherry::DRSH-1 co-lP’d with mCherry antibody or GFP antibody (recognizing PASH-1::GFP or PASH-1 (G-mut)::GFP). in, cell lysate input fraction; IP, co-IP fraction. Tubulin is shown as a loading control. 3 biological replicates are shown, except for wild type. (C) Western blot analysis of PASH-1::GFP and PASH-1(W-mut)::GFP co-lP’d with mCherry antibody (recognizing mCherry::DRSH-1) or GFP antibody. in, cell lysate input fraction; IP, co-IP fraction. Tubulin is shown as a loading control. 3 biological replicates are shown, except for wild type. (D) Western blot analysis of mCherry::DRSH-1 co-lP’d with mCherry antibody or GFP antibody (recognizing PASH-1 ::GFP or PASH-1(W-mut)::GFP). in, cell lysate input fraction; IP, co-IP fraction. Tubulin is shown as a loading control. 3 biological replicates are shown, except for wild type. (E-F) Western blot analysis of Pasha and Drosha interaction with FLAG::PASH-1(DD) in 2 independent experiments. Ratio of IP to input was 20:1 but note that in these blots, 2.5x protein equivelents were loaded for the G-mut and W-mut co-lPs to make them directly comparable to unaltered PASH-1::GFP. (E) Western blot analysis of FLAG::PASH-1(DD), PASH-1::GFP, and PASH-1 ::GFP(G-mut) co-lP’d with FLAG antibody (recognizing FLAG::PASH-1(DD)). in, cell lysate input fraction; IP, co-IP fraction. Tubulin is shown as a loading control. 2 biological replicates are shown, except for wild type and *mCherry::drsh-1 pash-1 ::GFP.* (F) Western blot analysis of FLAG::PASH-1(dd), PASH-1::GFP, PASH-1 ::GFP(G-mut), PASH-1 ::GFP(W-mut), and mCherry::DRSH-1 co-lP’d with FLAG antibody (recognizing FLAG::PASH-1(DD)). in, cell lysate input fraction; IP, co-IP fraction. Tubulin is shown as a loading control. 2 biological replicates are shown, except for wild type and *mCherry::drsh-1 pash-1 ::GFP,* which have only 1.

**Supplemental Figure S7.**
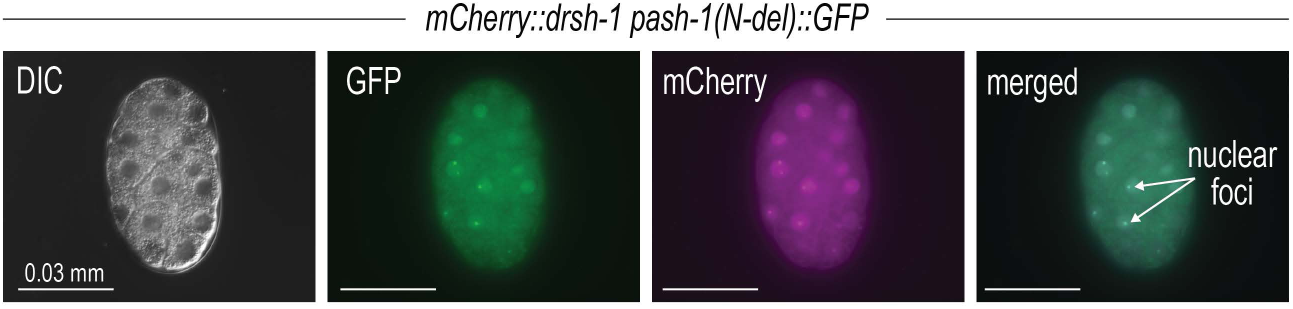
PASH-1(N-del)::GFP colocalizes with mCherry::DRSH-1. Representative images of PASH-1::GFP, PASH-1 (N-del)::GFP, and mCherry::DRSH-1 in embryos. DIC, GFP, mCherry, and GFP-mCherry merged images are shown.

